# Stony Coral Tissue Loss Disease biomarker bacteria identified in corals and overlying waters using a rapid field-based sequencing approach

**DOI:** 10.1101/2021.02.17.431614

**Authors:** Cynthia C. Becker, Marilyn Brandt, Carolyn A. Miller, Amy Apprill

## Abstract

Stony Coral Tissue Loss Disease (SCTLD) is a devastating disease. Since 2014, it has spread along the entire Florida Reef Tract, presumably via a water-borne vector, and into the greater Caribbean. It was first detected in the United States Virgin Islands (USVI) in January 2019. To more quickly identify disease biomarker microbes, we developed a rapid pipeline for microbiome sequencing. Over a span of 10 days we collected, processed, and sequenced coral tissue and near-coral seawater microbiomes from diseased and apparently healthy *Colpophyllia natans*, *Montastraea cavernosa*, *Meandrina meandrites* and *Orbicella franksi*. Analysis of the resulting bacterial and archaeal 16S ribosomal RNA sequences revealed 25 biomarker amplicon sequence variants (ASVs) enriched in diseased tissue. These biomarker ASVs were additionally recovered in near-coral seawater (within 5 cm of coral surface), a potential recruitment zone for pathogens. Phylogenetic analysis of the biomarker ASVs belonging to *Vibrio, Arcobacter,* Rhizobiaceae, and Rhodobacteraceae revealed relatedness to other coral disease-associated bacteria and lineages novel to corals. Additionally, four ASVs (*Algicola*, *Cohaesibacter*, *Thalassobius* and *Vibrio*) were exact sequence matches to microbes previously associated with SCTLD. This work represents the first rapid coral disease sequencing effort and identifies biomarkers of SCTLD that could be targets for future SCTLD research.

## 1. INTRODUCTION

Stony Coral Tissue Loss Disease (SCTLD) is a rapidly progressing, persistent, and widespread coral disease that affects at least 24 reef-building coral species in the Caribbean (Florida Keys National Marine Sanctuary, 2018). Since 2014, when it was first detected off Miami-Dade county, FL, it has devastated Floridian reefs, where loss in live coral has been as high as 60% (Precht *et al*., 2016; Walton *et al*., 2018). In the following years, the disease spread over the entire Florida Reef Tract and in 2018, the disease had appeared in disparate areas of the Caribbean (Kramer *et al*.). The SCTLD outbreak is one of the longest and most widespread coral disease outbreaks ever to be recorded. The extended duration, widespread occurrence, and high species susceptibility associated with SCTLD make this an unprecedented and devastating coral disease.

When SCTLD first emerged on Floridian reefs, it first impacted species including *Meandrina meandrites, Dichocoenia stokesii, Dendrogyra cylindrus* (an Endangered Species Act-listed coral), and the brain corals (i.e. *Colpophyllia natans, Pseudodiploria strigosa*). In ensuing months after the initial outbreak, other species began to show signs of SCTLD, including bouldering-type corals such as *Montastraea cavernosa,* and *Orbicella* spp. (Florida Keys National Marine Sanctuary, 2018). Similar assemblages of affected species and disease ecology confirmed the emergence of SCTLD on reefs off of Flat Cay, an unoccupied island off of St. Thomas, U.S. Virgin Islands (USVI), in January 2019. Tissue loss on corals in the USVI progress at rates up to 35-fold higher than other common coral diseases and leads to complete mortality of over half of afflicted colonies (Meiling *et al*., 2020). Throughout 2019, and until the time of sampling in February 2020, the disease spread around the island of St. Thomas and east to the island of St. John (Brandt *et al*., unpublished).

The pathogen or suite of pathogens responsible for SCTLD remain elusive, a common feature of the majority of coral diseases (Mera and Bourne, 2018; Vega Thurber *et al*., 2020). Successful cessation of lesion progression following application of amoxycillin paste to afflicted colonies suggests that either the pathogen(s) are of bacterial origin or that bacteria play a major role in disease progression and virulence as opportunistic microbes (Aeby *et al*., 2019; Neely *et al*., 2020). To detect putative pathogens or disease biomarker bacteria and archaea, 16S ribosomal RNA (rRNA) gene sequencing approaches that target bacteria and archaea were employed on field collected coral samples from the Florida Reef Tract, where the disease originated (Meyer *et al*., 2019; Rosales *et al*., 2020). Meyer and colleagues (2019) found bacteria from five genera, including *Vibrio, Arcobacter, Algicola, Planktotalea,* and one unclassified genus that were consistently enriched in the lesion tissue of three species of diseased corals (*Montastraea cavernosa*, *Diploria labyrinthiformis* and *Dichocoenia stokesii*). In a separate study (Rosales *et al*., 2020), Rhodobacterales and Rhizobiales sequences were associated with coral lesions in *Stephanocoenia intersepta*, *D. labyrinthiformis*, *D. stokesii* and *Meandrina meandrites*. It remains to be seen if these same microbial taxa are associated within SCTLD lesions across the greater Caribbean, especially in the geographically distant USVI, as well as in other coral species affected by the disease.

With such a widespread occurrence, it is important to understand the ways in which SCTLD is transmitted. It is hypothesized that transmission of this disease is through the water column, as evidenced by tank-based experiments (Aeby *et al*., 2019) and modeling of likely dead coral material and sediments within neutrally buoyant water parcels (Dobbelaere *et al*., 2020). Additionally, disease-associated microbial taxa were recovered in water and sediment of diseased-afflicted coral reefs, indicating that sediment may also play a role in transmission (Rosales *et al*., 2020). Disease-associated taxa or putative pathogens have yet to be examined in seawater directly surrounding diseased coral colonies using a targeted sampling method, such as syringe-based water sampling over corals (Weber *et al*., 2019).

The seawater directly overlying coral, here termed “near-coral seawater”, is an important reef environment. Compared to surrounding reef seawater, this environment is characterized by microbes with unique metabolisms and more virulent-like and surface-associated lifestyles. Also, this near-coral seawater environment is hypothesized to be a recruitment zone for both symbiotic microorganisms and potential pathogens (Weber *et al*., 2019). While coral physiology may influence the microbes living in this environment, water flow and surrounding currents also play a role (Silveira *et al*., 2017). Given existing evidence and hypotheses that the SCTLD pathogen or pathogens are water-borne (Aeby *et al*., 2019; Dobbelaere *et al*., 2020; Rosales *et al*., 2020), directly targeting the zone of potential pathogen recruitment is important for supporting these claims.

The rapid spread, persistence, and extreme virulence of SCTLD make it imperative to develop rapid response procedures for identifying candidate pathogens or biomarkers that could potentially be used in identifying the disease. Microbial community-based characterization approaches are ideal due to the breadth of information they provide, but can fall short in processing time. Typical microbiome pipeline procedures involve weeks to months of sample processing, sequencing, and data analysis, and this timeline is often impacted by other lab projects, personnel schedules and wait times at sequencing facilities. Furthermore, specialized equipment is often needed, making these procedures often impossible in remote island reef locations with limited laboratory support and facilities. In those instances, sample collection and shipment back to institutions with molecular laboratories and sequencing cores are the only options. Field-based sequencing circumvents these challenges and offers additional benefits, such as working immediately with fresh samples, and in the case of marine disease studies, the quickly processed data could even inform sampling strategies during the timeline of the project (Apprill, 2019). Recently, Illumina Inc. developed the iSeq 100 System, a portable sequencing platform. The platform uses sequencing by synthesis chemistry combined with complementary metal-oxide semiconductor (CMOS) technology and produces 8 million reads, with greater than 80% of reads passing a quality score of Q30 (99.9% base call accuracy) in each run (Illumina, Inc., 2020). Additionally, the CMOS technology allows the sequencing to occur all in a small, single-use cartridge, contributing to its ease of use in “pop-up” laboratory settings. More portable sequencing technology and the increasing availability of portable thermocycler machines and centrifuges have made it possible to set up molecular labs in almost any environment with an electrical connection.

Here, we developed and applied a rapid coral microbiome sequencing pipeline designed to more quickly gather data on the effects of SCTLD currently affecting numerous nations and reefs across the Caribbean. By setting up small, portable molecular biology tools in a home rental, we successfully collected, processed, and sequenced diseased and apparently healthy coral tissue and near-coral seawater samples at two reefs in St. Thomas, USVI (Fig. 1). We were interested in answering the following questions regarding the implementation of this rapid pipeline to more broadly understand the etiology of SCTLD: (1) How effective is a portable sequencing approach for coral disease studies, and potentially other marine diseases, (2) What microbial taxa are differentially distributed in healthy and diseased coral tissues, and which taxa biomarkers of the disease, (3) Can we identify SCTLD biomarker taxa in the seawater directly overlying healthy and diseased corals, and (4) To what extent are these SCTLD biomarkers phylogenetically related to known or unknown coral-associated bacteria?

**Fig. 1.**
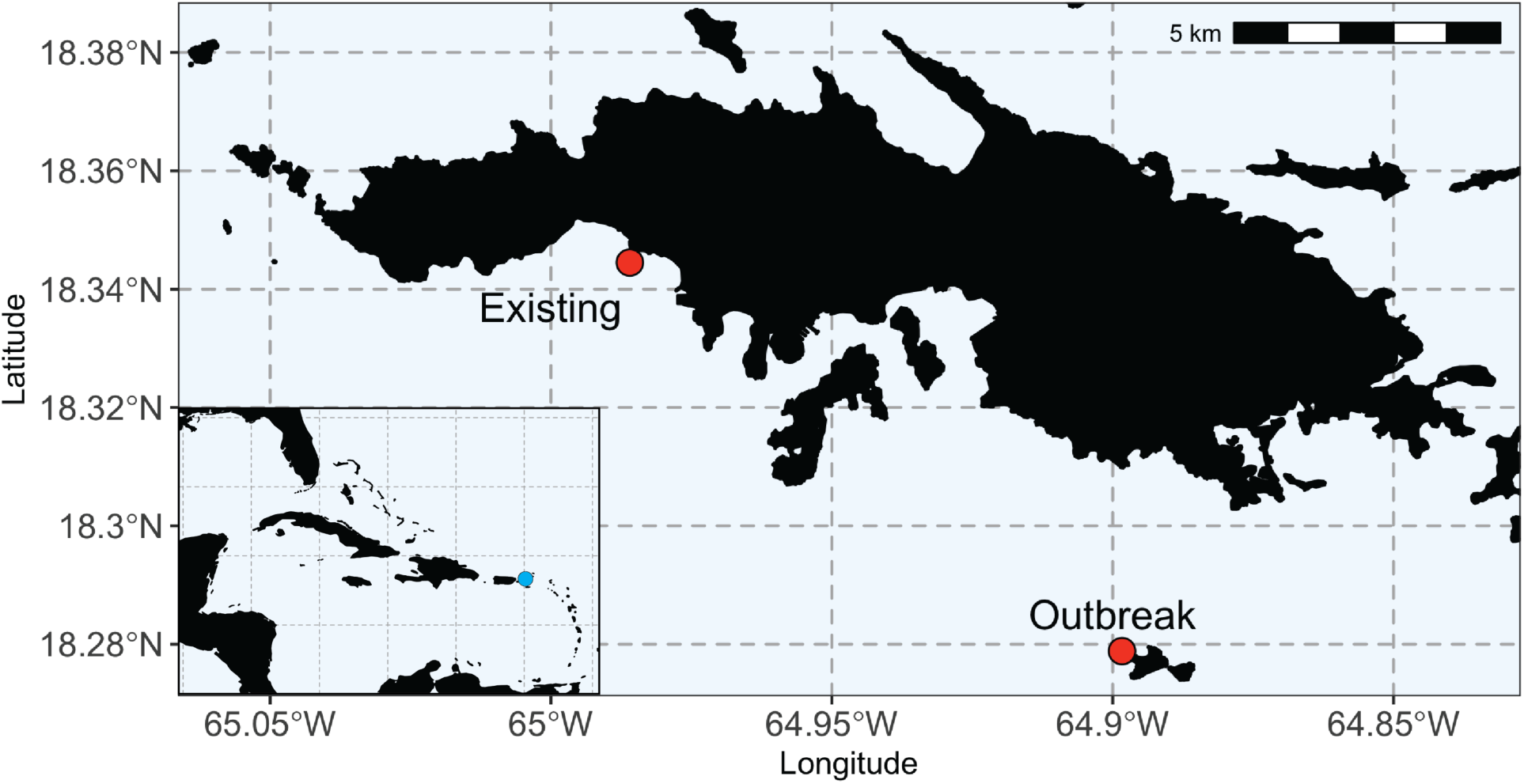
Sampling locations at Existing and Outbreak locations in St. Thomas, U.S. Virgin Islands. St. Thomas sampling locations included a reef at Black Point (Existing), which was experiencing SCTLD for 13 months prior to sampling and a reef at Buck Island (Outbreak), which had recently received SCTLD (less than one month affected). Scale bar is 5 km with marks at every kilometer. Inset map shows the greater Caribbean with the blue dot noting the location of the U.S. Virgin Islands.

**Fig. 2.**
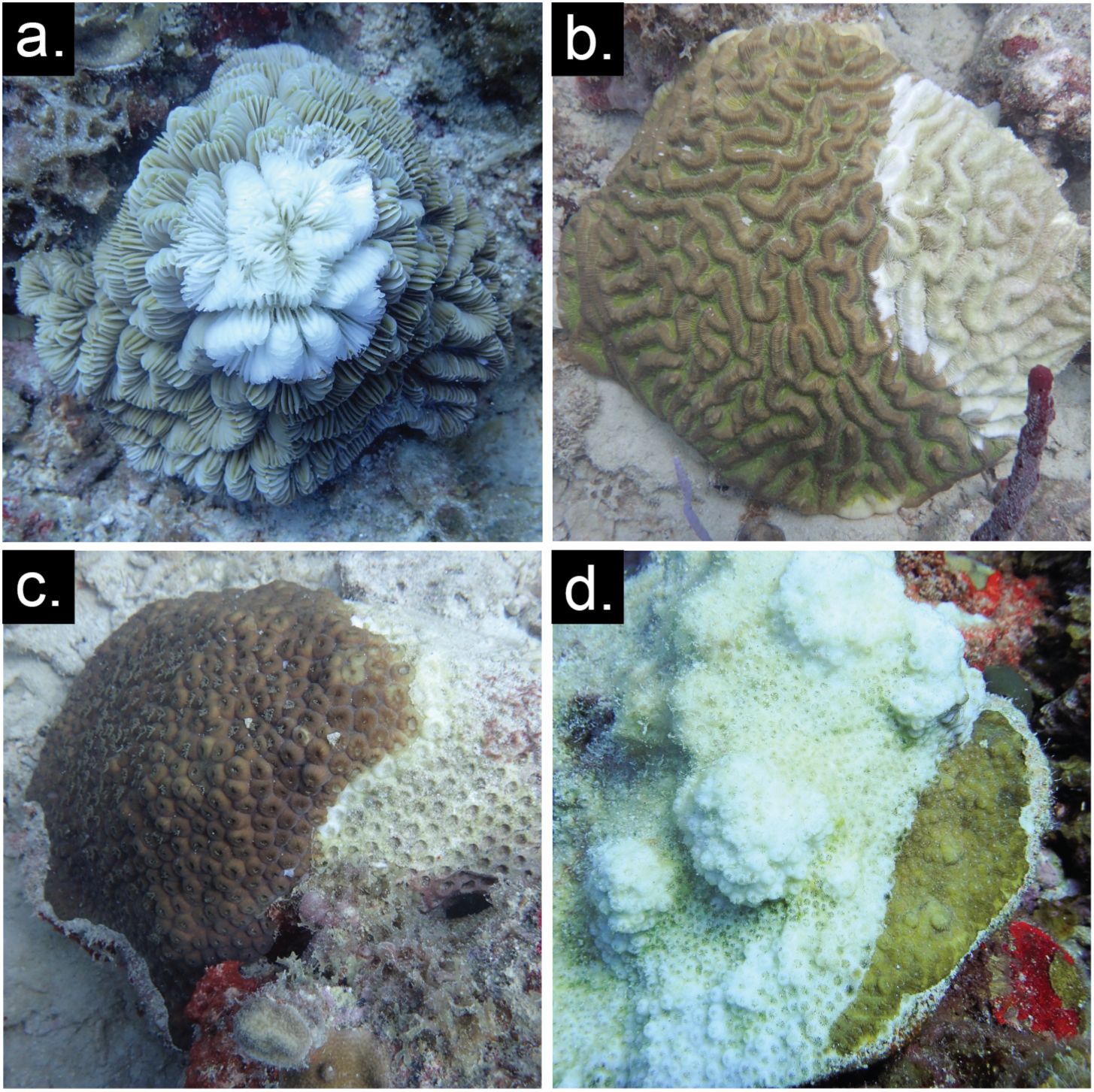
Stony Coral Tissue Loss Disease lesions progress across healthy tissue. Photos represent typical disease appearance on the following corals included in the present study: (a) *Meandrina meandrites,* (b) *Colpophyllia natans,* (c) *Montastraea cavernosa,* (d) *Orbicella franksi*. Seawater and tissue were sampled at the lesion front and 10 cm away from the lesion, or as far as possible from the lesion, when possible.

## 2. RESULTS

### 2.1 Output of field-based sequencing in portable microbiome laboratory

Three field-based sequencing runs with the Illumina iSeq 100 system each generated about 2 GB of paired-end, 150 bp sequencing data. In total, 12,997,634 sequencing reads were produced and used for subsequent data analysis. Each sequencing run lasted approximately 17 hours, and the runs were conducted on sequential days from February18th to 20th, 2020. The three days of sequencing produced high quality reads, with the forward reads containing 89.6%, 94.8%, and 89.8% of reads having a Q30 score or better, respectively. Following filtration of forward sequencing reads, the number of reads per sample for the 49 samples of coral tissue ranged from 60,105 to 128,036, with an average of 99,177 (± 14,511) while the range for the 51 seawater samples was 68,527 to 119,141, with an average of 96,933 (± 10,218) (Table 2). Thus, all samples were successfully sequenced with sufficient sequence reads for downstream analysis (60,000+ sequences). The average numbers of sequences recovered from the controls were as follows (average ± 1SD, n): Syringe Method Control samples (96,290 ± 7,542, n = 9), DNA extraction control samples (19,418 ± 11,672, n = 6), and PCR negative control samples (9,930 ± 904, n = 3) (Table 2). Over the course of the three sequencing runs, the same mock community of 20 known bacteria was sequenced to verify consistency and success of sequencing and ASV generation over the three individual runs. Successful identification of all 20 exact amplicon sequence variants within the mock community was achieved from all three runs, though for each, additional sequence variants were also recovered.

**Table 1.** Number of near-coral seawater (SW) and coral tissue (Coral) samples collected from the ‘Outbreak’ and ‘Existing’ SCTLD reefs on St. Thomas, USVI.

|  | Outbreak |  |  | Existing |  |  |
| --- | --- | --- | --- | --- | --- | --- |
|  | HH | HD | DD | HH | HD | DD |
| *Montastraea cavernosa* | 3 | 3 | 3 | 3 SW*<br>1 Coral* | 4 SW*<br>5 Coral* | 4 SW*<br>3 Coral* |
| *Colpophyllia natans* | 0 | 3 | 3 | 0 | 5 | 5 |
| *Meandrina meandrites* | 1 | 4 | 4 | 0 | 0 | 0 |
| *Orbicella franksi* | 0 | 3 | 3 | 0 | 0 | 0 |

\*Sample sizes from *M. cavernosa* from the Existing disease reef were different between seawater and coral due to sampling and processing constraints.

**Table 2.**
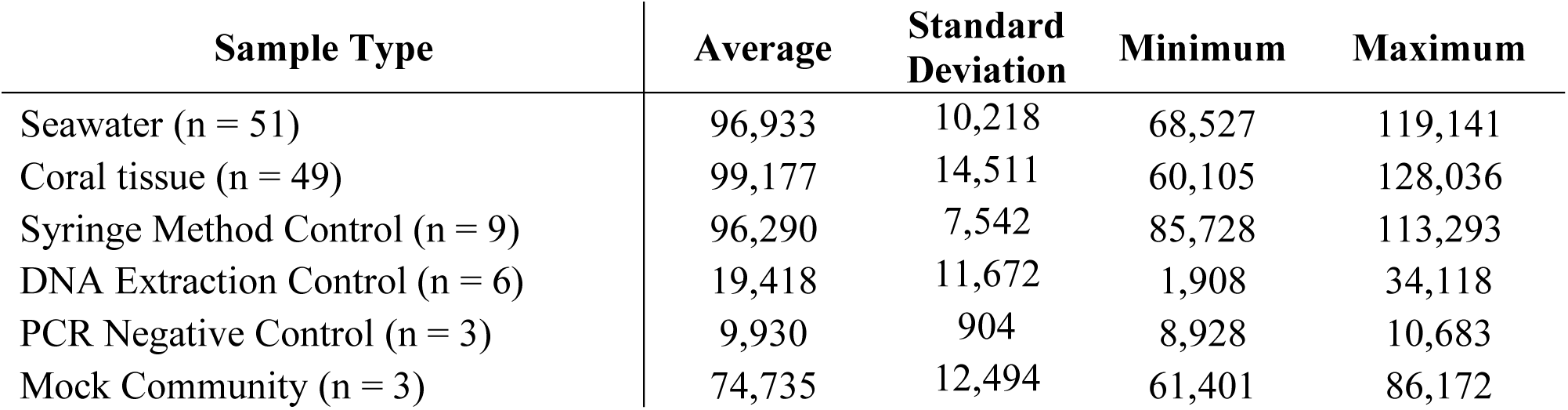
Summary statistics of sequencing reads produced by three sequencing runs on the Illumina iSeq 100 System, outlined by sample type.

### 2.2 Health state determines coral tissue microbiome structure while near-coral seawater microbiomes change according to site

Visualization of microbial beta diversity in coral tissue using Principal Coordinates analysis of Bray-Curtis dissimilarity revealed significant changes in the coral tissue microbiota associated with health condition, coral species, and reef site (Fig. 3a, Fig. S1-S2). Permutational Analysis of Variance (PERMANOVA) tests comparing diseased tissue (“DD”) to healthy tissues pooled from two sample types (healthy tissues from diseased colonies, “HD”, and healthy tissue from healthy colonies, “HH”; pooling occurred because tissue from apparently healthy colonies from all species was not available) revealed disease state as having a significant effect on coral tissue microbiome composition (Fig. 3a, PERMANOVA, R^2^ = 0.25, p < 0.001). The effect of health state, irrespective of species, was slightly higher when split up by all three conditions (“HH”, “HD”, “DD”, Fig. 3a, PERMANOVA, R^2^ = 0.26, p<0.001). The effect of species on structuring tissue microbial communities was also significant, though the effect size was smaller than between healthy and diseased coral tissue samples (Fig. 3a, Fig. S2, PERMANOVA, R^2^ = 0.15, p<0.001). Interestingly, a PERMANOVA where disease state was nested within coral species exerted an even greater effect, explaining 36% of microbiome structure (Fig. 3a, Fig. S2, PERMANOVA, R^2^ = 0.36, p<0.001). Together, disease state, species, and the nested designation exerted a larger effect on the microbial community composition compared to site-based changes, though site did significantly structure the coral tissue microbial communities (Fig. 3a, Fig. S2, PERMANOVA, R^2^ = 0.041, p = 0.031).

**Fig. 3.**
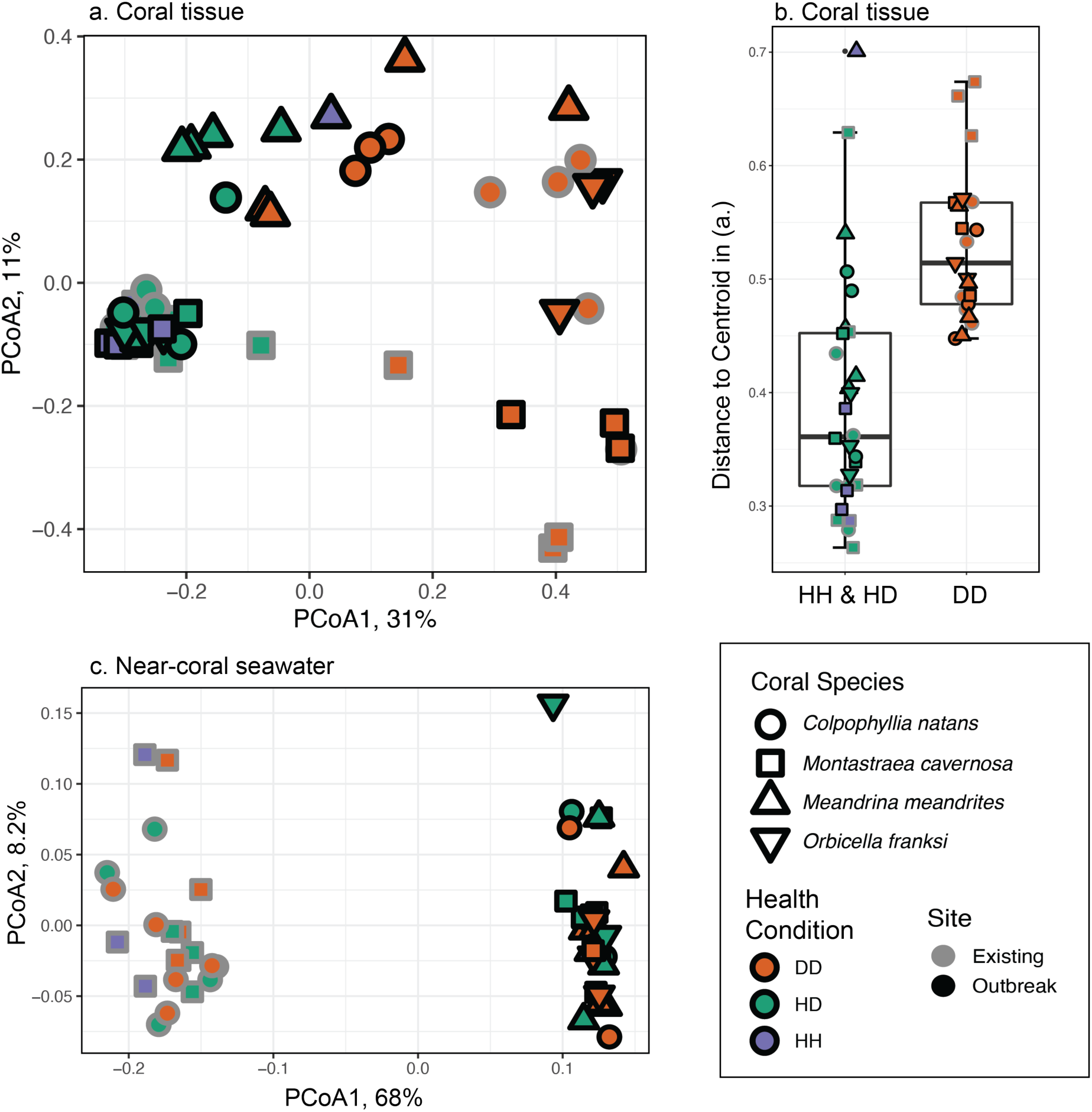
Coral tissue microbiomes differed according to health condition and near-coral seawater microbiomes differed according to site. (a) Principal coordinate analysis (PCoA) displays Bray-Curtis dissimilarity of coral tissue microbial communities, (b) beta diversity dispersion of coral tissue microbiomes represented by boxplots of the distance to centroid in (a), and (c) PCoA of Bray-Curtis dissimilarity in near-coral seawater microbiomes. Fill color represents health condition of diseased tissue (DD, orange), healthy tissue from a diseased colony (HD, green), or healthy tissue from an apparently healthy colony (HH, purple). Outline color indicates the reef where the sample was taken, which had either existing SCTLD infection (gray), or was experiencing a recent (<1 month) outbreak of SCTLD (black). Shape represents species of coral sampled: *Colpophyllia natans* (circle), *Montastraea cavernosa* (square), *Meandrina meandrites* (up triangle), and *Orbicella franksi* (down triangle).

Analysis of dispersion of beta diversity revealed significant differences between coral tissue health states (healthy vs. diseased) (Fig. 3b). The distance to centroid of all healthy tissue samples (HH and HD) was significantly lower, though more variable, than that of diseased samples (independent Mann-Whitney U Test, p < 0.001, Fig. 3b). Diseased tissue microbiome beta diversity dispersion was higher and more consistent compared to healthy tissue microbial beta diversity (Fig. 3b). Healthy tissue microbiomes were generally less dispersed (more closely clustered in the PCoA) except for a few samples, which were dispersed farther from the other healthy tissues (Fig. 3a,b). Furthermore, the range in raw Bray-Curtis dissimilarity values within each tissue sample type (healthy vs. diseased), reinforced the finding of increased beta diversity dispersion of diseased compared to healthy tissues (Mann-Whitney U Test p < 0.001, Fig. S3).

The effect of disease state was also visible in the stacked bar plot of coral tissue microbiomes (Fig. S4). Notably, *M. cavernosa* contained increased relative abundances of Deltaproteobacteria in diseased tissues (Fig. S4a). *Clostridia* and *Campyobacteria* relative abundances were increased in diseased tissues across all corals, though *Clostridia* was most prominent in diseased *M. cavernosa* and *C. natans*, while *Campylobacteria* was most prominent in diseased *O. franksi.* (Fig. S4). Interestingly, Oxyphotobacteria (predominantly *Prochlocococcus* and *Synechococcus*) decreased in relative abundance in diseased coral tissue (Fig. S4).

Near-coral seawater microbiomes taken within 5 cm of the coral surface were clearly distinct from tissue microbiomes, but structured according to site (PERMANOVA R^2^ = 0.67, p < 0.001) and not disease state (PERMANOVA R^2^ = 0.005, p = 0.921) (Fig. 3c, Fig. S1). Coral species also significantly affected the composition of the overlying seawater (Fig. 3c, PERMANOVA, R^2^ = 0.22, p = 0.004).

### 2.3 Disease biomarker bacteria identified within coral tissue

To detect specific bacteria that may be biomarkers of SCTLD, we used the beta-binomial regression model of the *corncob* R package (Martin *et al*., 2020) to test for differentially abundant ASVs between healthy (both HH and HD) and diseased (DD) tissue on each coral species. The model recovered 25 ASVs that were significantly more abundant, i.e. enriched and herein referred to as biomarkers, in the diseased tissue of at least one of the coral species (Table 3, Fig. 4, Fig. S5-S8). Ten of those 25 ASVs were enriched in diseased tissue of more than one coral species but none were enriched in all species (Table 3, Fig. S5-S8). Only ten of the 25 disease biomarker ASVs were significantly enriched in more than one coral species. Nonetheless, some ASVs, such as ASV44 (*Fusibacter*), were enriched in diseased coral tissue of all coral species, though the trend was not always significant. The 25 biomarker ASVs classified as belonging to 12 Families and 14 genera. Families with multiple biomarker ASVs were Arcobacteraceae, Desulfovibrionaceae, Family XII of the order Clostridiales, Rhodobacteraceae, and Vibrionaceae (Table 3). Within these, four ASVs belonged to the genera *Arcobacter*, five to *Vibrio*, and three to *Fusibacter*.

**Fig. 4.**
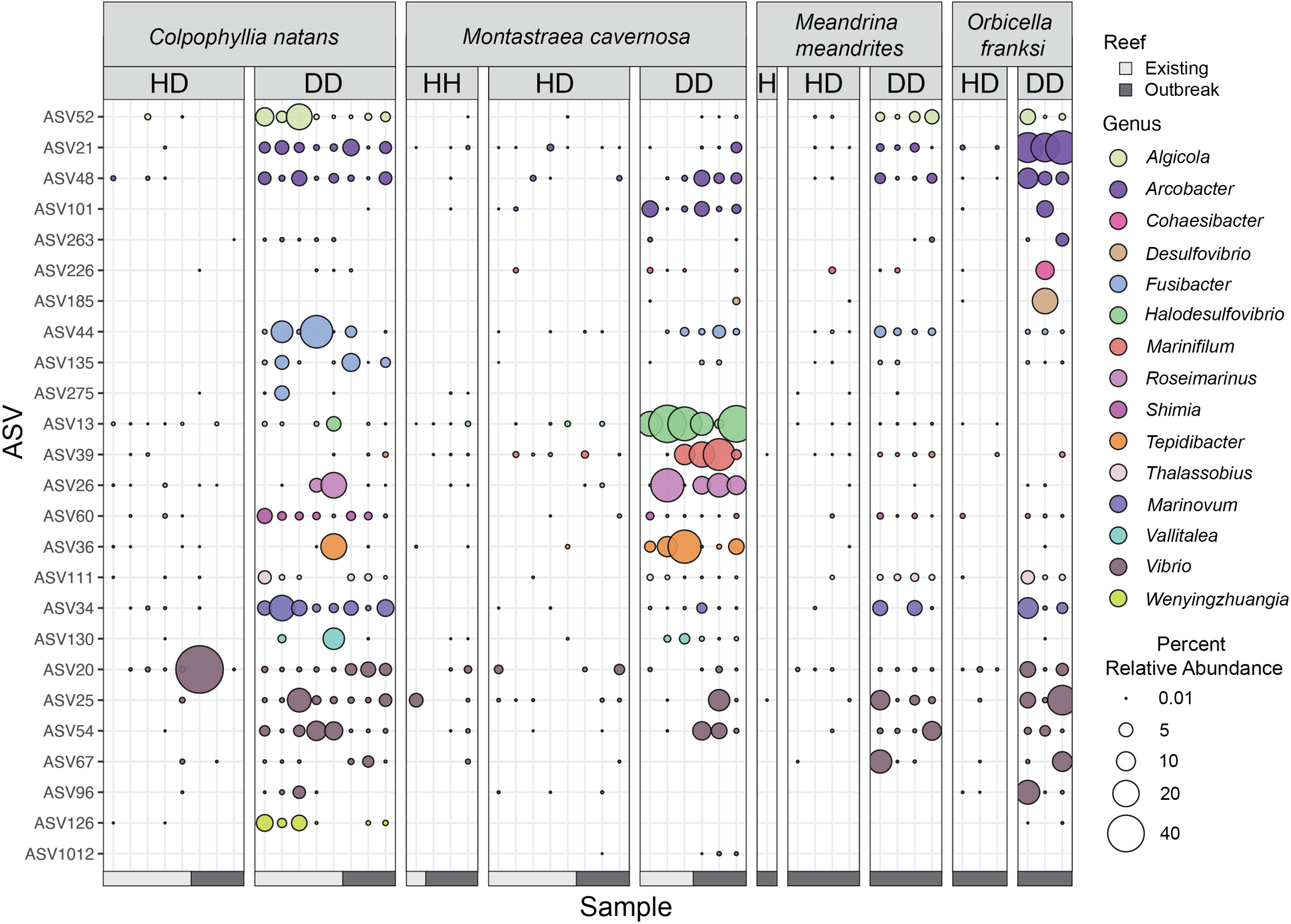
Relative abundances of 25 SCTLD biomarker ASVs significantly differentially enriched (FDR corrected p-value < 0.05) in diseased tissue of at least one coral speices. Samples on the x-axis are organized by coral species (*Colpophyllia natans*, *Montastraea cavernosa*, *Meandrina meandrites*, and *Orbicella franksi*), health state of the coral (healthy tissue from apparently healthy colony = “HH”, healthy tissue from diseased colony = “HD”, disease lesion tissue = “DD”). Additionally, a color bar at the bottom indicates the coral was collected at the Existing disease reef (light gray) or at the disease Outbreak reef (dark gray). ASVs on the y-axis are organized and colored by Genus. Percent relative abundance of each ASV is represented by the size of the colored circle, with a percent relative abundance of zero represented by the absence of a circle or dot. The relative abundances were calculated after removing common seawater bacteria and archaea, which were determined using the syringe method control samples containing ambient reef seawater and with the R-package *decontam* (see methods).

**Table 3.**
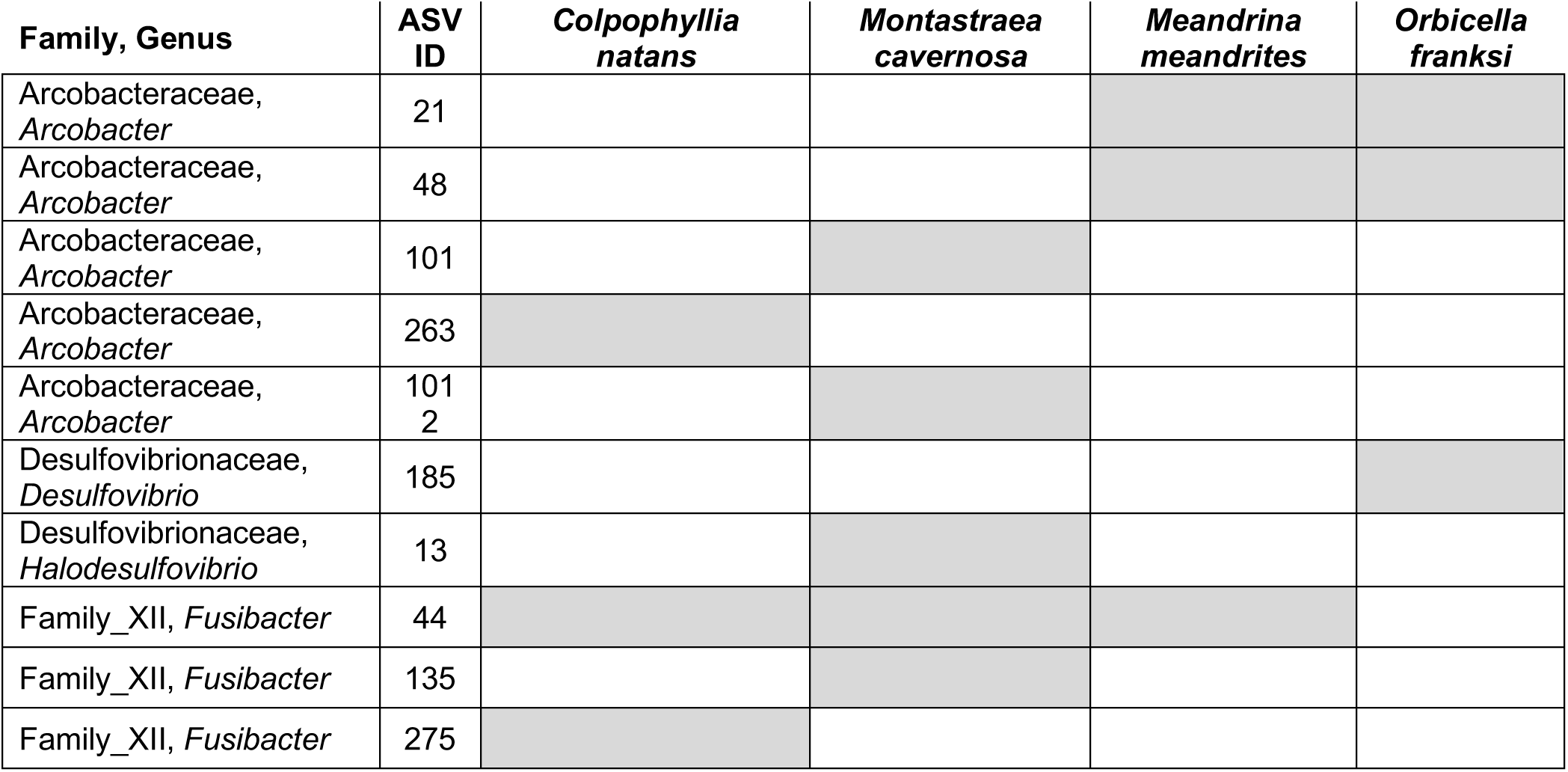
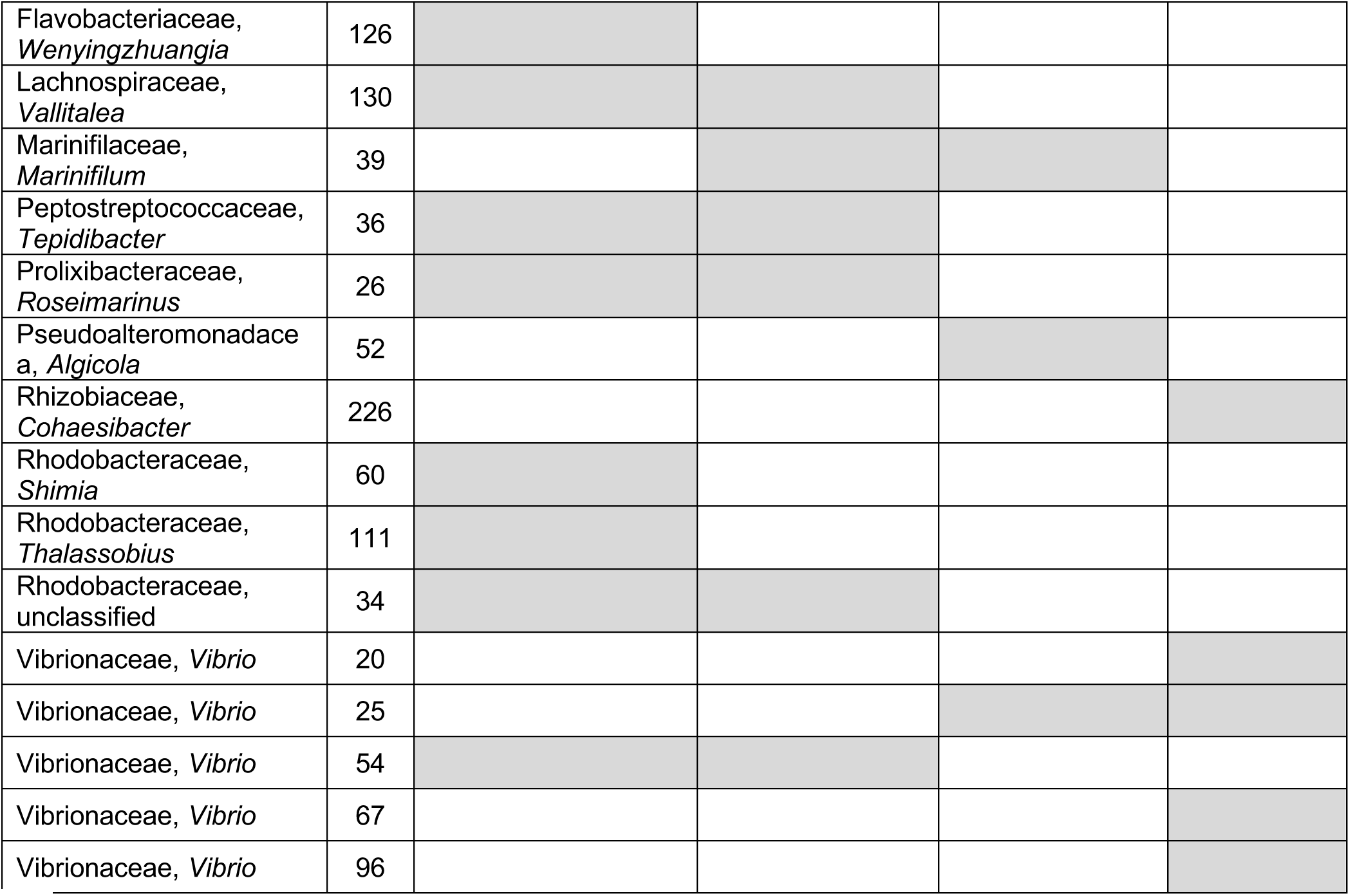
Summary of coral species featuring significant enrichment (FDR corrected p-value < 0.05) of SCTLD biomarker ASVs in disease lesion tissue (DD) compared to healthy tissue (HH and HD combined). White cells indicate no significant difference in the relative abundance of the ASV between healthy and diseased tissue. Differential abundance of ASVs was calculated using the beta-binomial regression model of the R-package *corncob* and ASVs were considered significant at an FDR corrected p-value <0.05.

In addition to identifying ASVs enriched in diseased tissue, the differential abundance analysis revealed other ASVs that were depleted in diseased tissue relative to healthy tissue, and were therefore healthy tissue-associated (coefficient < 0, Fig. S5-S8). *M. meandrites* tissue had only one ASV enriched in healthy tissue (Family Terasakiellaceae from Rhodospirillales; Fig. S7). Healthy tissue of *C. natans* (Fig. S5) and *M. cavernosa* (Fig. S6) were enriched with ASVs belonging to Clades Ia, Ib, and unclassified Clade II of SAR11, the NS4 Marine Group and NS5 Marine Group of Flavobacteriaceae, *Prochlorococcus* MIT9313*, Synechcococcus* CC9902, and unclassified SAR116. Healthy tissue of *C. natans* also was enriched in ASVs belonging to unclassified Rhodobacteraceae, Clade 1a Lachnospiraceae, and OM60 (NOR5) clade of Halieaceae. Unique healthy tissue-associated ASVs in *M. cavernosa* included an unclassified Clade IV of SAR11, NS2b Marine Group of Flavobacteriaceae, *Candidatus* Actinomarina, unclassified genera of the AEGEAN-169 Marine Group, Urania-1B-19 marine sediment group of Phycisphaeraceae, unclassified Cryomorphaceae, *Coraliomargarita*, unclassified Ectothiorhodospiraceae, R76-B128 Kiritimatiellaceae, unclassified Pirellulaceae, and *Blastocatella* (Fig. S6). Healthy tissue-associated ASVs from *O. franksi* largely represented taxa found in healthy tissue from at least one of the other three coral species targeted in this study (Fig. S8). One ASV that classified as an unclassified Arcobacteraceae was associated with *O. franksi* healthy tissue, as well as two ASVs that classified to the genus *Endozoicomonas* (ASV99, ASV108).

### 2.4 Disease biomarker taxa recovered within near-coral seawater

Previous studies indicated that seawater may be a vector for the SCTLD pathogen or pathogens; therefore, we hypothesized that the seawater within 5 cm of coral lesions would harbor the 25 SCTLD biomarker ASVs we identified from coral tissue. A differential abundance test of the 25 biomarker ASVs in seawater overlying disease lesions (DD) compared to healthy tissue (HH and HD) did not find significant enrichment of those ASVs in seawater over diseased lesions. We further tested the 25 biomarker ASVs in near-coral seawater over apparently healthy colonies compared to disease lesion colonies of *M. cavernosa,* the only species for which we had sufficient replication of apparently healthy colonies. Again, there was no ASV significantly enriched in waters overlying diseased corals. Despite the lack of significant enrichment of putative pathogens within near-coral lesion seawater, we did observe all 25 biomarker ASVs in the near-coral seawater over diseased tissues, except for two ASVs (*Cohaesibacter* ASV226 and *Desulfovibrio* ASV185). Several of the SCTLD-associated ASVs were present in seawater overlying all four diseased coral species, including *Algicola* (ASV52), *Arcobacter* (ASV21, ASV101), *Halodesulfovibrio* (ASV13), *Marinifilum* (ASV39), and *Vibrio* (ASV20) (Fig. 5). Interestingly, ASV34, an unclassified Rhodobacteraceae, was found only in near-coral seawater directly overlying disease lesions, but not over healthy tissue across all species (Fig. 5). Overall, disease-enriched bacteria were identified at low levels (<1.5% relative abundance) in near-coral seawater, though there was no significant enrichment of these taxa over diseased tissue.

**Fig. 5.**
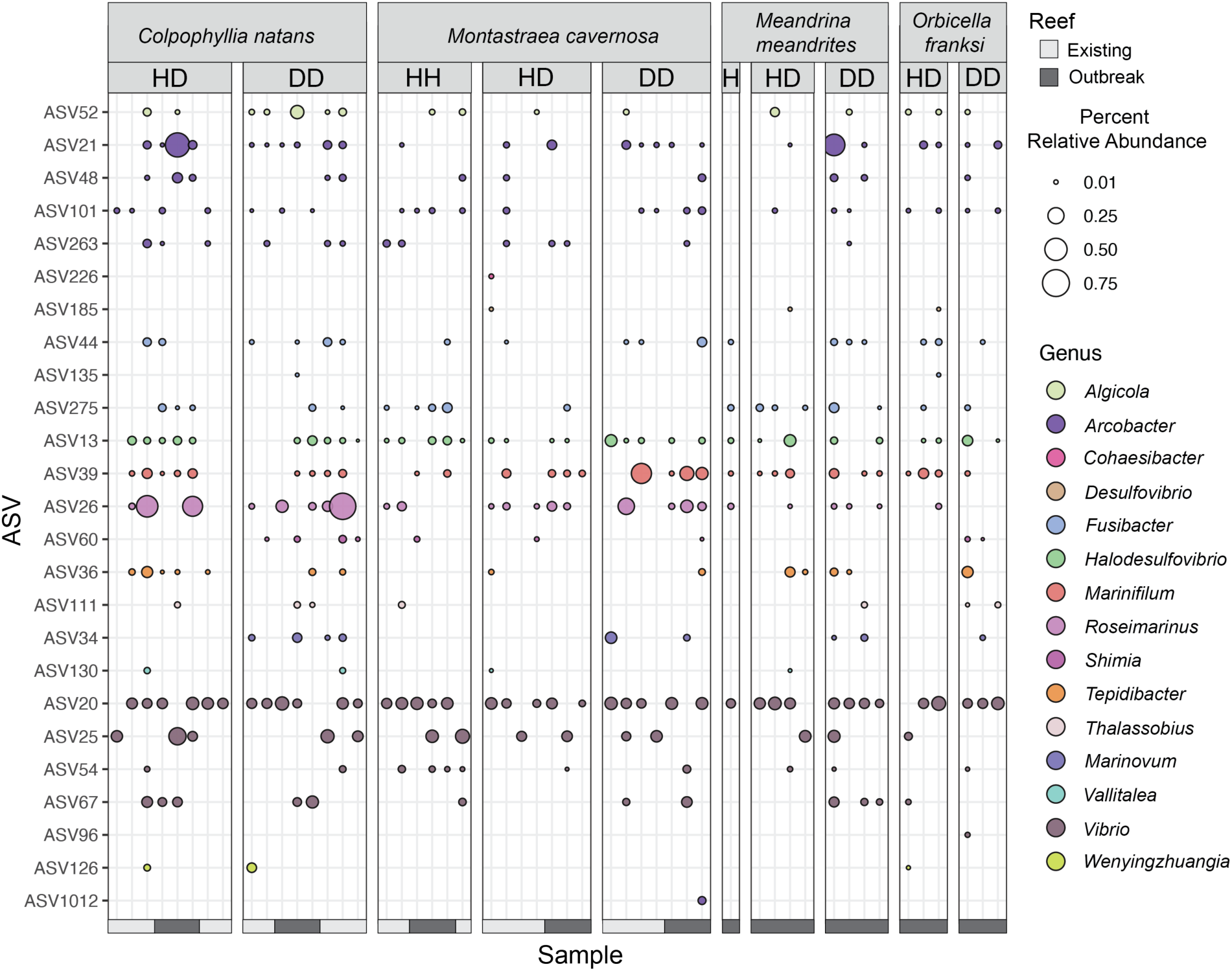
SCTLD biomarker ASVs identified in coral tissue found in near-coral seawater. Samples on the x-axis are organized by coral species (*Colpophyllia natans*, *Montastraea cavernosa*, *Meandrina meandrites*, and *Orbicella franksi*), and health state of the coral (healthy tissue from apparently healthy colony = “HH”, healthy tissue from diseased colony = “HD”, disease lesion tissue = “DD”). Additionally, a color bar at the bottom indicates whether the coral was collected at the Existing disease reef (light gray) or at the disease Outbreak reef (dark gray). ASVs on the y-axis are organized and colored by Genus. Percent relative abundance of each ASV is represented by the size of the colored circle, with a percent relative abundance of zero represented by the absence of a circle or dot. ASVs graphed are those identified by differential abundance analysis as significantly enriched in diseased coral tissue (FDR corrected p-value < 0.05) of at least one coral species.

### 2.5 Phylogenetic analysis of SCTLD-enriched taxa

Given the high representation of *Arcobacter*, *Vibrio,* Rhizobiaceae and Rhodobacteraceae ASVs in the biomarker ASVs, we produced phylogenetic trees to better predict species-level identifications and to relate the ASVs to other sequences associated with corals and coral diseases. Phylogenetic analysis of the Campylobacterota (formerly Epsilonbacteraeota) genus *Arcobacter* spp. indicated no close association of the SCTLD-associated ASVs to *Acrobacter* isolates, but did detect close association of ASV101 and ASV48 with several clone sequences from Black Band Disease in *Siderastrea siderea* and other corals (Fig. 6). Similarly, ASV21 was related to a sequence recovered from an unidentified coral reef disease, while ASV263 was most closely related to a previously identified coral-associated sequence (Fig. 6). Overall, all SCTLD-associated *Arcobacter* spp. ASVs from the current study grouped in clades with coral-associated or coral disease sequences (Fig. 6).

**Fig. 6.**
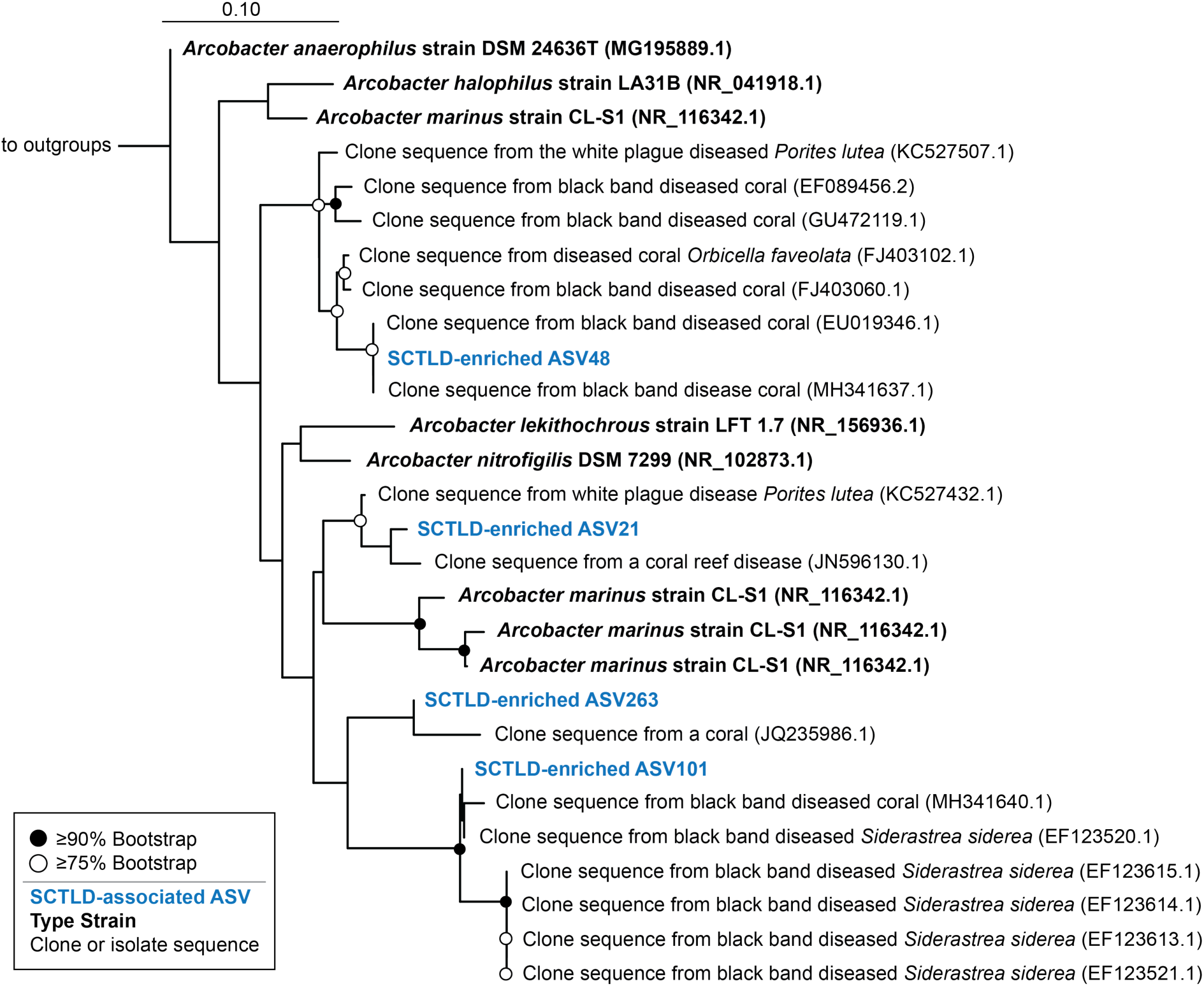
Biomarker ASVs from the genus *Arcobacter* closely related to isolates and clone sequences from diseased corals. Reference phylogenetic tree was produced using RAxML rapid bootstrapping with an automatic bootstrapping approach to produce the highest-scoring maximum likelihood tree using only longer-length sequences (black). SCTLD-associated ASVs (blue) identified by differential abundance analysis or by previous studies were added to the tree using the Evolutionary Placement Algorithm in RAxML. Colors represent qualitative information about the sequences as follows: Blue = SCTLD-associated ASVs from the present study, black bold = bacterial type strains, black = clone or bacterial isolate/strain sequences. GenBank accession numbers are located in parentheses following each taxa label. Circles at node represent bootstrap values of ≥ 90% (filled-in circle) or ≥ 75% (empty circle). Tree was rooted using the 16S rRNA gene of *Streptococcus mutans* strain ATCC 25175 (NR_115733.1).

Phylogenetic analysis of the SCTLD biomarker ASVs from the gammaproteobacterial genus *Vibrio* spp. leveraged a reference tree previously constructed using existing coral-associated sequences found in the Coral Microbiome Database (Huggett and Apprill, 2019) and *Vibrio* type strains. Additionally, given the previous identification of a SCTLD-associated *Vibrio* ASV by Meyer and colleagues (2019), we included that sequence in this phylogenetic tree. ASV20 closely aligned to *V. harveyi* ATCC 35084, an isolate obtained from a brown shark kidney following a mortality event (formerly known as *V. carchariae* (Grimes *et al*., 1984; Pedersen *et al*., 1998) (Fig. 7). Interestingly, *Vibrio* ASV54 in the present study was an exact sequence match to the SCTLD-associated *Vibrio* ASV reported previously (Meyer *et al*., 2019), and this sequence is novel to corals (Table 4, Fig. 7). Biomarker ASV96 aligned near Vibrio Cluster (VC) 24 and near an isolate from a cold-water coral (Fig. 7). ASV67 was closely related to both *Vibrio pectenicida* and a sequence recovered from a white plague-afflicted coral (Fig. 7). Similar to the other ASVs, ASV25 was related to other coral-associated *Vibrio* sequences. Overall, most *Vibrio* biomarker ASVs were identified as unique coral-associated sequences.

**Fig. 7.**
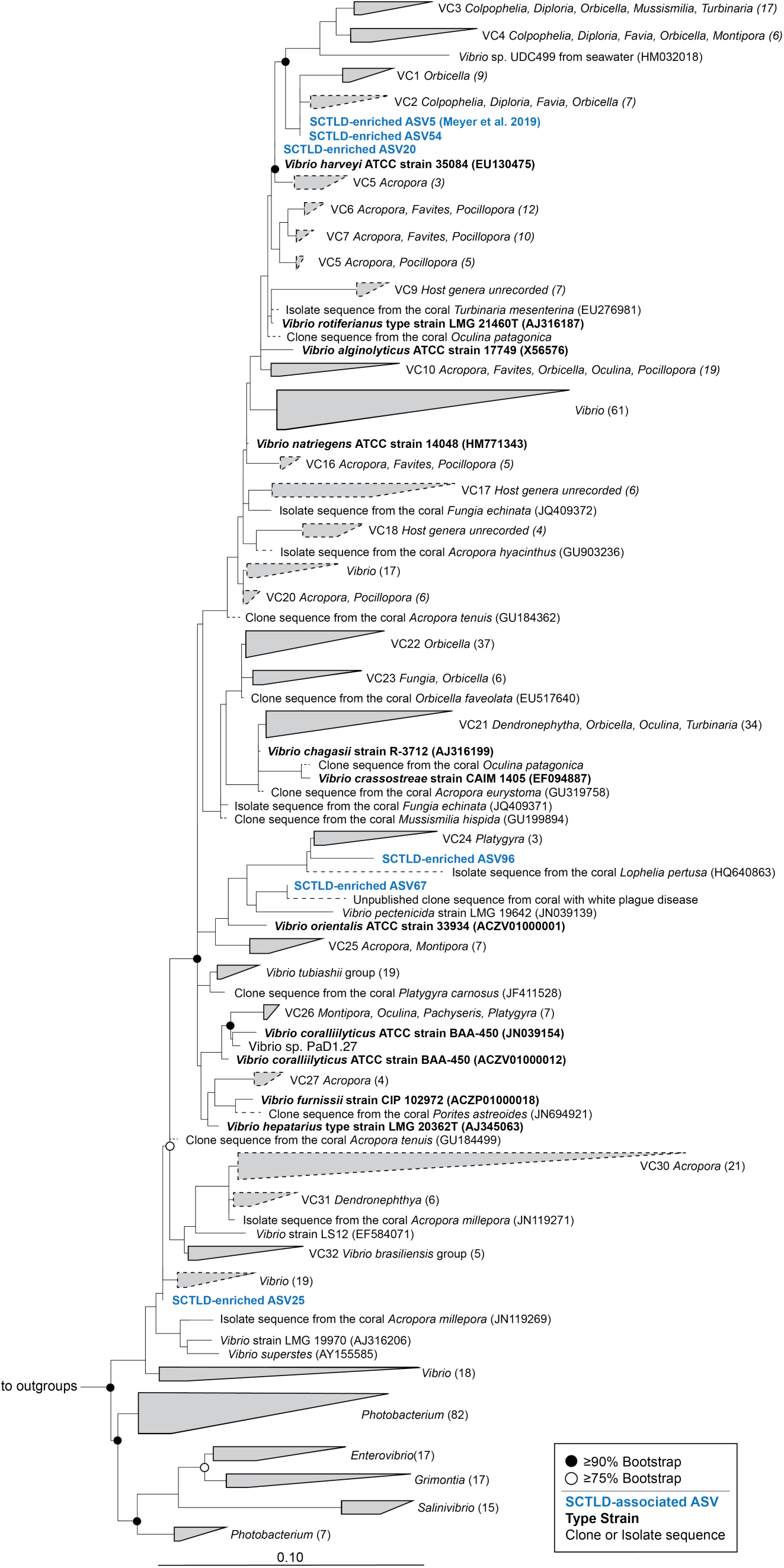
Biomarker *Vibrio* ASVs from the present study and a recent study related to *Vibrio* pathogens, type strains, and sequences obtained from the Coral Microbiome Database. Maximum likelihood and bootstrapped phylogenetic tree was produced using RAxML based on long (>1200 bp) sequences only, with the shorter coral associated sequences (dashed lines) and SCLTD-associated sequences (blue text) added using the Quick-add Parsimony tool in ARB. Colors represent qualitative information about the sequences as follows: Blue = SCTLD- associated ASVs from the present or previous study (Meyer *et al*., 2019), Black bold = bacterial type strains, Black = clone or bacterial isolate/strain sequences. GenBank accession numbers are located in parentheses following each taxa label, when available. Circles at node represent bootstrap values of ≥ 90% (filled-in circle) or ≥ 75% (empty circle). Tree was rooted with *Thalassospira xianhensis* (EU017546) and *Thalassospira tepidiphila* (AB265822).

**Table 4.** Biomarker ASVs in the present study with 100% sequence similarity to SCTLD- associated ASVs identified by previous studies.

| Family | Genus | ASV ID in present study | Enriched in diseased tissue (Rosales et al. 2020) | Enriched in diseased tissue (Meyer et al. 2019) |
| --- | --- | --- | --- | --- |
| Pseudoalteromonadaceae | *Algicola* | 52 | no | **Yes** |
| Rhizobiaceae | *Cohaesibacter* | 226 | **Yes** | no |
| Rhodobacteraceae | *Thalassobius* | 111 | **Yes** | no |
| Vibrionaceae | *Vibrio* | 54 | no | **Yes** |

We produced a phylogenetic tree with all known SCTLD-associated ASVs that classified to Rhizobiaceae, sequences from other coral disease studies, and other related sequences (Fig. 8). One biomarker ASV from the present study (ASV226) was identical to a previous SCTLD- associated ASV11394 (Rosales *et al*., 2020), and both grouped with *Cohaesibacter marisflavi*, a bacterium that has been isolated from seawater (Table 4, Fig. 8). ASV18209 and ASV19474 also fell within the *Cohaesibacter* genus, and were most closely aligned to sequences isolated from White Plague affected corals. The *Pseudovibrio* SCTLD-associated ASV19959, ASV30828, and ASV16110 from Rosales *et al*. (2020) were most closely related to *Pseudovibrio denitrificans* NRBC 100300 compared to other *Pseudovibrio* type strains (Fig. 8). Finally, the unclassified Rhizobiaceae ASV34211 (Rosales *et al*., 2020) was most closely associated to isolates of *Hoeflea* spp. and ASV24311 (Rosales *et al*., 2020) to *Filomicrobium* spp. (Fig. 8).

**Fig. 8.**
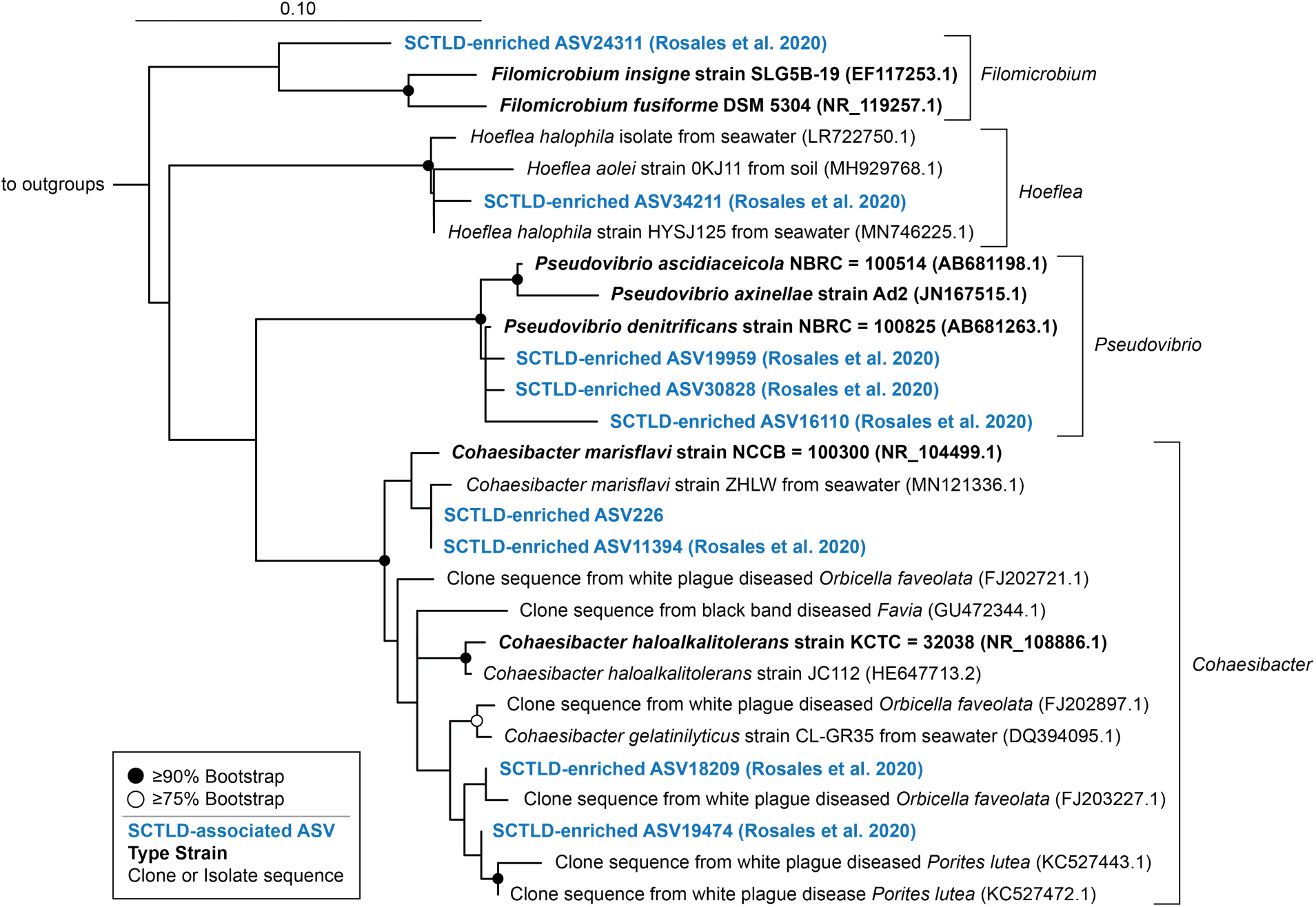
One SCTLD biomarker Rhizobiaceae ASV from the present study and several from a previous study related to other Rhizobiaceae sequences associated with corals and coral diseases. Reference phylogenetic tree was produced using RAxML rapid bootstrapping with an automatic bootstrapping approach to produce the highest-scoring maximum likelihood tree using only longer-length sequences (black). SCTLD-associated ASVs (blue) identified by differential abundance analysis or in a previous study (Rosales *et al*., 2020) were added to the tree using the Evolutionary Placement Algorithm in RAxML. Colors represent qualitative information about the sequences as follows: Blue = SCTLD-associated ASVs from the present or a previous study (Rosales *et al*., 2020), Black bold = bacterial type strains, Black = clone or bacterial isolate/strain sequences. GenBank accession numbers are located in parentheses following each taxa label. Circles at node represent bootstrap values of ≥ 90% (filled-in circle) or ≥ 75% (empty circle). Tree was rooted using the 16S rRNA gene of *Streptococcus mutans* strain ATCC 25175 (NR_115733.1).

Phylogenetic analysis of SCTLD biomarker ASVs that classified as Rhodobacteraceae using a reference tree generated from the Coral Microbiome Database (Huggett and Apprill, 2019) revealed close classification to bacteria associated with coral hosts that were distinct from existing isolates (Fig. 9). Several SCTLD-associated ASVs from Rosales and colleagues (2020) (ASV15252, ASV24736, ASV13497, ASV3538, and ASV29944) were related to sequences from ballast water and hypersaline mats rather than coral sequences (Fig. 9). In contrast, the SCTLD biomarker ASV60 from the present study was matched to other coral-associated *Rhodobacteraceae* sequences with no definitive classification, though related to *Phaeobacter* (Fig. 9). Several *Rhodobacteraceae* ASVs classified to *Thalassococcus* sequences (ASV29894, ASV25482, ASV29283 from Rosales *et al*. (2020)) and ASV111 from the present study (exact sequence match to ASV29283 from Rosales *et al*. (2020)). Finally, SCTLD-associated ASV34 classified within the likely *Marinovum* genus alongside many coral-associated sequences (Fig. 9).

**Fig. 9.**
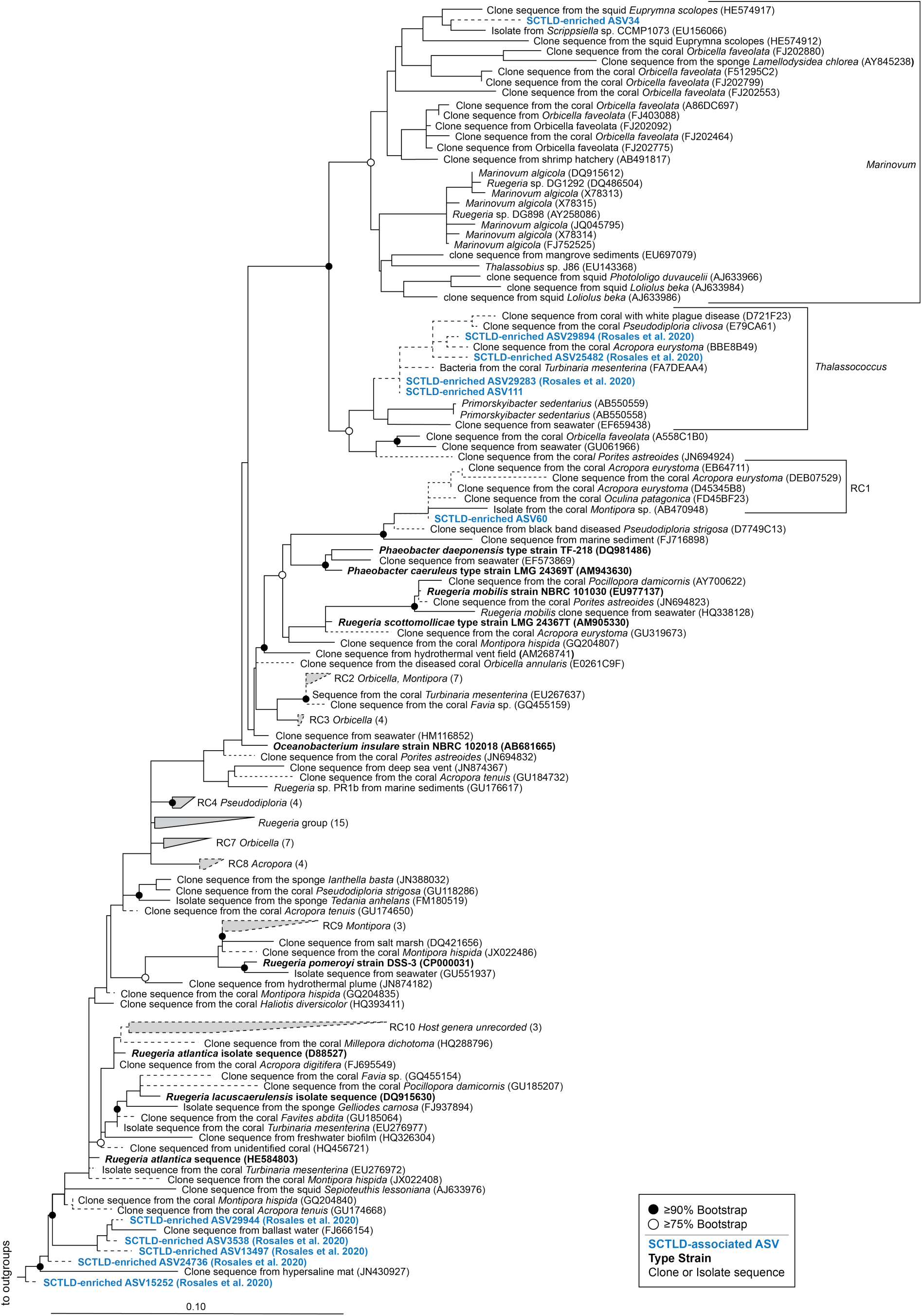
Two SCTLD biomarker Rhodobacteraceae ASVs from the present study and several from a previous study related to sequences from the Coral Microbiome Database encompassing several genera within the Rhodobacteraceae Family. Maximum likelihood and bootstrapped phylogenetic tree was produced using RAxML based on long (>1200 bp) sequences only, with the shorter coral associated sequences (dashed lines) and SCLTD-associated sequences (blue text) added using the Quick-add Parsimony tool in ARB. Colors represent qualitative information about the sequences as follows: Blue = SCTLD-associated ASVs from the present or a previous study (Rosales *et al*., 2020), Black bold = bacterial type strains, Black = clone or bacterial isolate/strain sequences. GenBank accession numbers are located in parentheses following each taxa label, when available. Circles at node represent bootstrap values of ≥ 90% (filled-in circle) or ≥ 75% (empty circle). Tree was rooted with *Alteromonas* (AACY023784545) and *Methylophilaceae* (HM856564 and EU795249).

## 3. DISCUSSION

To better understand the effect of Stony Coral Tissue Loss Disease on coral reefs in the U.S. Virgin Islands, an area with an active and detrimental SCTLD outbreak (VI-CDAC), we developed and integrated a rapid, field-based 16S rRNA gene sequencing approach to characterize microbiomes of coral tissue and near-coral seawater of SCTLD-infected colonies. This is the first study on SCTLD microbiomes from the U.S. Virgin Islands. Additionally, this represents the first application of the Illumina iSeq 100 System to coral disease. St. Thomas, USVI does not have a molecular ecology laboratory or sequencing facility; therefore, we transformed a home rental on the island to a molecular laboratory, and in the span of two weeks, we carried out a complete microbiome workflow, from sample collection to sequencing. This short timeline enabled us to process fresh samples, gather data more quickly, and begin data analysis in the following months, which revealed significant differences between healthy and diseased coral tissue, regardless of coral species or reef location. Differential abundance analysis identified 25 SCTLD biomarker ASVs, all of which were present in the seawater directly overlying coral. Furthermore, prominent biomarker ASVs represented sequences highly related to the *Vibrio harveyi/V. carchariae* pathogen, sequences unique to corals, and many ASVs phylogenetically related to bacterial sequences previously identified in diseased corals.

### 3.1. SCTLD lesion microbial communities are unique from healthy tissue communities

We identified clear and consistent differences between healthy and diseased coral microbiomes, regardless of location and species of coral. In addition, the dispersion of beta-diversity was consistently higher among the diseased corals, compared to all healthy tissue samples, which had reduced, yet more variable dispersion of beta diversity. This greater beta diversity across diseased coral tissue microbiomes could be related to disease etiology or methodological factors. The syringe-based collection method results in homogenized seawater, skeleton, tissue, and mucus collected together at the disease lesion interface. As a result, the disease sample may include apparently healthy tissue, newly compromised, diseased tissue, as well as necrotic or sloughed off tissue. Thus, this mixture likely captures potential pathogen(s), organisms involved in secondary infections, or even saprophytic microorganisms proliferating off the exposed skeleton and dead coral tissue (Burge *et al*., 2013; Egan and Gardiner, 2016). For example, the finding of increased Deltaproteobacteria in diseased tissues of *M. cavernosa,* and the significant enrichment of *Halodesulfovibrio,* known sulfate-reducing bacteria, may have been a signature of the exposed coral skeleton (Chen *et al*., 2019), or perhaps anaerobic degradation of coral tissue (Viehman *et al*., 2006). Overall, the finding that disease impacts coral microbiome structure in the USVI is supported by previous findings that show shifts in coral microbiomes between healthy and diseased coral tissues in Florida, USA (Meyer *et al*., 2019; Rosales *et al*., 2020).

Differential abundance analysis between healthy and diseased tissue microbial communities revealed 25 disease biomarker ASVs, which may represent potential pathogens or opportunistic bacteria. Of these, we identified one known pathogen (ASV20), with 100% similarity in the overlapping region to *V. harveyi* (formerly *V. carchariae*; EU130475.1) isolated from a shark mortality event, and shown to be virulent for spiny dogfish (*Squalus acanthias*)(Grimes *et al*., 1984). This ASV was consistently detected in diseased coral tissue (DD samples), including all *C. natans, M. meandrites* and *O. franksi* colonies, and two-thirds of the *M. cavernosa* colonies, at relative sequence abundances of 5% or lower. Additionally, this ASV was frequently recovered in the HD and HH colonies, and most near-coral seawater samples, indicating its broad prevalence in these diseased reefs. Interestingly, *V. harveyi* has been suggested as the causative agent of white syndrome disease in aquaria and field-based corals (Luna *et al*., 2010) and is also associated with diseases in flounder and other fish, sharks, abalone, shrimp and sea cucumbers (Austin and Zhang, 2006). Despite the prevalence of *V. harveyi* sequences in the present SCTLD study, this was not a SCTLD-associated bacterium identified in the Florida-based studies. Still, it seems relevant to examine pathogenicity of *V. harveyi* in future coral disease experiments.

Four of our SCTLD biomarker ASVs were identical to ASVs identified in Florida-based SCTLD studies. *Algicola* ASV52 and *Vibrio* ASV54 were identical to SCTLD-associated ASVs identified by Meyer *et al*. (2019) and both sequences were also recovered in other Caribbean coral diseases (black band disease or white plague disease type II; reviewed by Meyer *et al*. 2019). In addition to association with disease, phylogenetic analysis placed ASV54 and the previously-identified ASV5 (Meyer *et al*., 2019) as phylogenetically distinct from other known coral-associated lineages, suggesting it may be an invasive bacterium (Fig. 7). Although the not significant in all corals in this study, the noticeable enrichment in abundance of both ASV52 (*Algicola*) and ASV54 (*Vibrio*) in all corals may be biologically relevant, given the association of *Algicola* and *Vibrio* in SCTLD-affected corals sampled in Florida approximately 1,800 km away (Meyer *et al*., 2019).

Two other biomarker ASVs in the present study, *Cohaesibacter* (ASV226) and *Thalassobius* (ASV111), were also enriched in another study of Florida-based SCTLD-associated microbiomes by Rosales et al. (2020). In our study, *Cohaesibacter* ASV226 was enriched in diseased tissue of only *O. franksi,* and was found at low relative abundances or not at all in the rest of the corals. Rosales *et al*. (2020) detected the same *Cohaesibacter* ASV in *S. intercepta, D. labyrinthiformis,* and *M. meandrites* affected by SCTLD. Phylogenetic analysis placed those two similar sequences (ASV226 and ASV11394, Fig. 8) with *Cohaesibacter marisflavi*, a species not currently known to be a pathogen (Qu *et al*., 2011). Although no *Cohaesibacter* species are known pathogens, *Cohaesibacter intestini* was isolated from the intestine of an invertebrate, abalone (Liu *et al*., 2019). Two *Cohaesibacter* sequences identified by Rosales *et al*. (2020) were phylogenetically related to clone sequences isolated from white plague disease type II affected *Orbicella* (formerly *Montastraea*) *faveolata* (Sunagawa *et al*., 2009) and white plague affected *Porites lutea* (Roder *et al*., 2014). We therefore predict a role of novel *Cohaesibacter* species in coral disease that could be targets of future isolation studies. Additionally, *Thalassobius* (ASV111) was an exact match to one of eight Rhodobacteraceae ASVs enriched in diseased corals in the Rosales *et al*. (2020) study. In the present study, this ASV was significantly enriched only in *C. natans,* but was generally present in diseased tissue of all species of coral examined. Furthermore, these ASVs, and two other disease-associated ASVs (ASV29894 and ASV25482) from Rosales *et al*. (2020) classified to unique coral bacteria and sequences in the *Thalassococcus* genus, which included one sequence from a white plague disease afflicted coral. Interestingly, several of the SCTLD-associated ASVs added into the phylogenetic tree were completely distinct from previously reported coral-associated sequences and isolates, and instead represented sequences more closely associated to sequences from hypersaline mat or ballast water environments. Despite variability in putative identity of the diverse Rhodobacteraceae sequences associated with SCTLD, exact sequences recovered from diseased corals across geographic regions in the Caribbean (Florida, USA, and USVI) may indicate some concordance in the effect of this disease on different coral species regardless of geography.

It should be noted that there are some methodological differences between the SCTLD studies, which could impact the microbial sequences recovered and compared. The studies all utilized the same primers, but the sequencing platforms differed in read length; the previous studies used merged reads, enabling a total read length of approximately 253 bp, whereas this study only used 126 bp forward reads due to sequencing of primers. A different primer set targeting a smaller region would be necessary for reads to be merged. Otherwise, the three studies employed the DADA2 analysis pipeline, resulting in sequences published alongside ASV identifiers, allowing for comparison of amplicon sequence variants across studies, a significant benefit of the DADA2 pipeline (Callahan *et al*., 2017). Lastly, we did take care to insert the shorter amplicon sequences into a phylogenetic framework based on longer read sequences. While the placement of the ASVs appear robust, additional marker genes or genomes are necessary to confirm the taxonomies affiliated with the ASV-based sequences.

In addition to *V. harveyi* ASV20, the other 24 biomarker bacteria may also be worth investigating from the perspective of polymicrobial infections. Black band disease, which is commonly spotted on Caribbean reefs, is well-established as a polymicrobial disease. Black band disease results from interactions among cyanobacteria, sulfur-cycling bacteria, archaea, and even eukaryotic microorganisms that form a mat which migrates across the coral colony, killing tissues (Sato *et al*., 2016). Another common group of Caribbean coral diseases are white plague-type diseases. SCTLD was initially identified as a white plague-like disease, due to the similar presentation of a white expanding lesion that progresses across the coral (Precht *et al*., 2016). However, the fast progression, species affected, and unique ecology clearly differentiate it from other white plague diseases (Florida Keys National Marine Sanctuary, 2018; Meiling *et al*., 2020; Muller *et al*., 2020). White plague-like diseases have had bacterial pathogens identified, such as *Aurantimonas coralicida*, but often they are inconsistent or unidentified across studies (Denner, 2003; Sunagawa *et al*., 2009). Although SCTLD is unique from white plague-like and black band diseases, several SCTLD biomarker ASV sequences were phylogenetically similar to existing environmental sequences and isolates from white plague disease or black band disease-afflicted corals. This potentially indicates common opportunistic taxa that colonize diseased coral hosts, or may suggest a polymicrobial disease etiology for SCTLD.

Similar to previous reports for white plague disease, it could be that both bacteria and viruses play a role in SCTLD onset and virulence. Antibiotic pastes containing amoxicillin have been shown to be effective at slowing and halting progression of SCTLD (Aeby *et al*., 2019). While these results indicate bacterial involvement in SCTLD infection and virulence, viral infection may also play a role in this disease; a group of single-stranded DNA viruses have been shown to play a role in white plague-like diseases (Soffer *et al*., 2014). We did not investigate viruses in our study but metagenomic and microscopic techniques that investigate holobiont components, such as bacteria, archaea, DNA and RNA-based viruses, and fungi, should be employed in the future to further identify the etiology of this devastating and destructive disease. Additionally, a drawback of the current study is the lack of replication of samples from apparently healthy colonies (“HH”). While collection of apparently healthy colony tissue was limited in the present study, future work could aim to prioritize collection of tissue from apparently healthy colonies, as it would serve as an important baseline for comparison to samples from diseased or compromised hosts.

### 3.2. Signals of SCTLD infection in near-coral seawater

Biomarker bacteria identified as SCTLD-enriched were broadly recoverable in near-coral seawater (<5 cm) surrounding the coral colonies, though seven ASVs were found in fewer than five of the nine samples. The seawater less than 5 cm from the coral surface is within the momentum boundary layer, which features reduced flow due to friction of seawater over the organisms compared to seawater one meter and higher above the benthos (Shashar *et al*., 1996; Barott and Rohwer, 2012). Increased abundances of copiotrophic-type microbial lineages in near-coral seawater has led to a hypothesis that the region may be a recruitment zone for symbionts or potential pathogens (Silveira *et al*., 2017; Weber *et al*., 2019). The SCTLD biomarker taxa found within the coral momentum boundary layer, or near-coral seawater habitat, support this hypothesis. These potentially opportunistic or pathogenic taxa may have been responding to exuded DOM released from the coral host (Haas *et al*., 2011, 2013; Nelson *et al*., 2011, 2013). Indeed, coral species was a driver of microbiome structure in the near-coral seawater while disease state was not, supporting the idea that seawater microbial communities may have been responding to species-specific chemical signals (Nelson *et al*., 2013; Weber *et al*., 2019).

Interestingly, according to the differential abundance comparison, none of the disease biomarker ASVs were significantly enriched in seawater overlying diseased compared to healthy areas of the corals. A further test between near-coral seawater of apparently healthy colonies of *M. cavernosa* compared to disease lesions on *M. cavernosa* also failed to distinguish biomarker taxa at higher relative abundances in seawater over the diseased lesions. However, relative abundances may not be useful for identifying pathogens in seawater; information on pathogen load needed to cause and/or reflect diseased conditions may be more useful. Additionally, within near-coral seawater, the microbial communities are reflective of water movement (Silveira *et al*., 2017). In many cases, the distance between healthy and diseased near-coral seawater samples in our study was only 10 cm. In the time it took to sample each location, the seawater likely moved around and across the healthy and diseased sections of the colony, potentially causing a more homogeneous signal and obscuring any differences near-coral seawater microbiomes of diseased versus healthy tissue. Also, we did not include seawater from non-diseased reefs, which could be useful for deciphering pathogen signals in the future.

Beyond coral-based influences, the greatest driver of near-coral seawater microbial beta diversity in our study was reef location, though these changes were subtle compared to the shifts in coral tissue microbiome beta diversity. The influence of environmental shifts and location-based changes in seawater microbiomes was noted previously, and was even used to predict microbiome composition (Glasl *et al*., 2019). The two reefs we sampled were only approximately 12 km away from each other and featured quite similar environmental conditions (Table S1). While the conditions were quite similar, minor differences in these conditions may have caused minor differences in microbiome composition in the near-coral seawater; although, with only two locations, it is difficult to determine if, or which, environmental conditions were structuring the near-coral seawater microbial communities. Hydrographic conditions may also have played a role in structuring the seawater communities. The ‘Outbreak’ site was located offshore and likely experienced stronger currents and water flow than the ‘Existing’ disease location, which was partially enclosed in a relatively calm bay (Fig. 1).

While physicochemical conditions were only slightly different across reefs, a major distinction between the two reef locations was degree of coral disease and coral cover, which also may have played a role in structuring the near-coral seawater microbial community. Underlying differences in coral reef benthic composition can exert an influence on the taxonomic composition of reef seawater microbial communities (Kelly *et al*., 2014). At the time of sampling, coral cover at the ‘Existing’ site was still higher than at the ‘Outbreak’ site (11.5% ± 1.8 SEM compared with 6.0% ± 1.2 SEM, respectively), even though it had been affected by the disease for at least 13 months compared with only 1 month at the ‘Outbreak’ site. However, the ‘Outbreak’ site had nearly double the amount of disease prevalence (8.3% ± 4.0 SEM) compared with the ‘Existing’ site (4.3% ± 4.3 SEM) (Ennis *et al*., unpublished).

While seawater is hypothesized to be the disease vector (Aeby *et al*., 2019), our investigation failed to identify significantly greater abundances of disease-associated bacteria surrounding lesions compared to healthy tissue. Although we found largely homogeneous microbiome structure in the near-coral seawater, we still propose that sampling within this zone increases the likelihood of detecting the potentially water-borne pathogens compared to sampling ambient, or surrounding seawater outside of this zone. Furthermore, perhaps increased sample sizes targeting completely healthy colonies compared to diseased colonies, rather than seawater overlying healthy tissue on a diseased colony would allow for better identification of SCTLD-associated bacteria in seawater. Beyond seawater, recent evidence suggests that sediments surrounding coral may play an important role as a reservoir of SCTLD pathogens (Rosales *et al*., 2020) though that was not sampled here. Future investigations into SCTLD vectors should aim to sample both near-coral sediments and seawater, both *in situ* and in isolated mesocosm tanks to provide further information on the likely modes of transmission of SCTLD pathogens.

### 3.3. Rapid and portable microbiome profiling is feasible and applicable to marine diseases

Here we successfully implemented an in-the-field microbiome protocol to rapidly assess the shifts in microbiome composition associated with the destructive coral disease, SCTLD. The emergence of smaller, more portable DNA sequencing technology is critical for increasing accessibility to sequencing technology and curbing the long turnaround time of months-to-years required to process and sequence field-collected samples (Krehenwinkel *et al*., 2019). In the present study, we developed and applied a more portable protocol because of those attractive benefits of smaller sequencing technology. St. Thomas, USVI, does not currently contain a laboratory with the capacity for DNA sequencing. To efficiently conduct microbiome work, we set up our own laboratory. Furthermore, SCTLD is a fast-acting disease (Meiling *et al*., 2020), making a speedy collection-to-data turnaround time critical for distributing results and findings to reef managers more quickly and efficiently.

Smaller, portable sequencing technology began with the launch of the minION (Oxford Nanopore Technologies, Oxford, UK) in 2014. Since then, it has revolutionized and facilitated increased field-based sequencing efforts aimed at biodiversity assessments and other monitoring programs (reviewed by Krehenwinkel *et al*., 2019; Maestri *et al*., 2019). The minION sequencer has been effective at identifying fungal pathogens of plants (Hu *et al*., 2019), viral pathogens (Hoenen *et al*., 2016; Batovska *et al*., 2017), and antibiotic resistance in bacterial pathogens (Leggett *et al*., 2020). While long-read metagenome sequencing works well in many systems for pathogen identification, high levels of coral host and *Symbiodinaceae* algal symbiont DNA would have dominated long reads on the minION system. Instead, we pursued short read, amplicon-based microbiome sequencing on Illumina-based system, which allowed for the selective enrichment and sequencing of bacterial and archaeal sequences.

Illumina launched the iSeq 100 System only recently, in 2018. It is the smallest (1 foot cube) and most portable Illumina technology to date and features a single-use cartridge that houses all sequencing reagents, further contributing to its ease of use. The iSeq 100 System is built upon the sequencing-by-synthesis and short-read technology, and has a maximum read length of 150 bp. While this is significantly shorter than the read lengths possible by the minION (2-98 kb, (Laver *et al*., 2015)), the 150 bp sequences are sufficient for broad metabarcoding investigations and ASV generation. Given the need for Polymerase Chain Reaction (PCR) to amplify the barcode region, the pipeline we conducted was greatly aided by miniaturized PCR machines, including miniPCR 8 and the BentoLab, which were simple to use and easier to pack than standard benchtop thermocyclers. Additionally, smaller centrifuges and the small Qubit 2.0 fluorometer were easily packed and adapted for our home-rental laboratory. Overall, the rapid microbiome pipeline employed here performed well. Following three sequencing runs, the number of reads generated by the iSeq per sample was comparable to those recovered in a previous study of SCTLD microbiomes that used MiSeq sequencing for the same region of DNA and the same sample collection method (Meyer *et al*., 2019). Given these comparable read counts per sample and high-quality nature of the sequencing runs (89.6 - 94.8% of reads passing Q30), the Illumina iSeq 100 System could be an ideal target for future studies on marine microbial communities, especially in cases when disease outbreaks occur and there is a need for rapid information and results to better inform remediation and management of such disease outbreaks.

The present workflow could be applied again to SCTLD research. With all data analysis scripts saved and easily accessible on GitHub, future data could be easily processed and compared to the present findings during future in-the-field sequencing projects. This would contribute to the aspect of our pipeline that could have been improved, which is the length of time needed to process data, examine trends, and report results. While we had all of our data in 10 days after the project start, there was still significant time needed for investment into the data analysis component, which occurred back at our home institution and was intermixed with needs from other projects. Future work could focus on producing additional scripts that incorporate predictive, machine-learning algorithms to analyze the microbial communities in coral tissue and identify microbial predictors of SCTLD, similar to work that identified microbial predictors of environmental features within reef seawater microbiomes (Glasl *et al*., 2019). This could allow scientists the potential to identify corals afflicted with SCTLD before entire colonies are killed, and within the timeline of fieldwork or research cruises. Additionally, as more is learned about the identity of individual marine pathogens, then targeted pathogen identification approaches in novel systems may become more straightforward.

### 3.4. Conclusion

Stony Coral Tissue Loss Disease has collectively affected hundreds of kilometers of coastal and offshore reefs in the Caribbean, with no present indication of stopping. This study aimed to develop and implement a field-based, rapid microbiome characterization pipeline in the USVI, an area more recently affected by the SCTLD outbreak. Following successful sequencing on the Illumina iSeq 100, we identified 25 SCTLD biomarker ASVs that may represent putative pathogens, including, *V. harveyi*, a bacterium known to be pathogenic in other marine systems. Many of the 25 biomarker ASV sequences enriched in diseased tissue were recovered in near-coral seawater, a potential recruitment zone for pathogens and the hypothesized vector for SCTLD. Interestingly, four of the SCTLD biomarker ASVs identified in our study exactly match sequences previously reported as enriched in SCTLD lesion tissue. Phylogenetic analysis revealed that many of the disease biomarker ASVs were related to likely novel coral or coral-associated disease bacteria. Future investigations aimed at isolating and characterizing those microorganisms and other SCTLD biomarker bacteria would be able to better determine if these organisms are pathogens or opportunists, and how they potentially target and grow around or within coral hosts. In the present study, the successful integration of a rapid pipeline for studying coral disease generated data more quickly, and subsequent analysis revealed differences in microbiome structure associated with the SCTLD outbreak in the USVI. This contributes to the growing body of literature on SCTLD that is largely focused in Florida, USA. Finally, we found that this rapid microbiome characterization approach worked well for identifying microbial indicators of coral disease, and it may have useful applications to marine diseases more broadly.

## 4. EXPERIMENTAL PROCEDURES

### 4.1 Sample collection

Coral colonies showing active Stony Coral Tissue Loss Disease (SCTLD) and nearby completely healthy colonies were targeted for sampling on February 11 and 13, 2020 on Buck Island (18.27883°, −64.89833°), and Black Point (18.3445°, −64.98595°) reefs, respectively, in St. Thomas, USVI (Fig. 1). Buck Island was considered a recent outbreak site where disease first emerged in January 2020 and will be referred to as “Outbreak”, whereas Black Point had been experiencing SCTLD since at least January 2019, and will be referred to as “Existing”. Coral species sampled were *Montastraea cavernosa* (Outbreak and Existing)*, Colpophyllia natans* (Outbreak and Existing)*, Meandrina meandrites* (Outbreak), and *Orbicella franksi* (Outbreak; Table 1). SCTLD was identified by single or multi-focal lesions of bleached or necrotic tissue with epiphytic algae colonizing the recently dead and exposed skeleton (Fig. 2). At both reefs, some paling of colonies was apparent, especially on *Orbicella* spp., as a result of a recent bleaching event in October 2019. Due to this, it was challenging to distinguish SCTLD from white plague-type diseases, which generally occur following bleaching events (Miller *et al*., 2009). As a result, we avoided sampling *Orbicella* spp., except when it was clear the colony had regained full coloration and the disease lesion was consistent with SCTLD infection.

To investigate if putative pathogens were recoverable from seawater surrounding diseased colonies, near-coral seawater was sampled 2-5 cm away from each coral colony prior to tissue sampling via negative pressure with a 60 ml Luer-lock syringe (BD, Franklin Lakes, NJ, USA). Two seawater samples were collected over each colony displaying SCTLD lesions: one sample was taken directly above healthy tissue approximately 10 cm away from the lesion, when possible, and a second sample over diseased tissue. Syringes were placed in a dive collection bag for the duration of the dive. Once on board the boat, the seawater was filtered through a 0.22 μm filter (25 mm, Supor, Pall, Port Washington, NY, USA) and the filter with holder was placed in a Whirl-pak bag and kept on ice until returning to the shore. While on shore, filters were placed in sterile 2 ml cryovials (Simport, Beloeil, QC, Canada) and frozen in a liquid nitrogen dry shipper.

After near-coral seawater sampling, samples of tissue and mucus mixed together (hereafter referred to as ‘slurry’ samples) were collected. One slurry sample was collected from each healthy colony and two from each diseased colony. For the two samples collected from each diseased colony, one was collected from the interface between healthy and newly bleached tissue (Fig. 2), and the other from healthy tissue approximately 10 cm away from the disease interface. When limited healthy tissue remained on a diseased colony, the slurry was collected approximately 3 to 5 cm away from the disease lesion interface. The tissue and mucus slurries were collected with 10 ml non-Luer lock syringes (BD) by agitating and disrupting a small area of the tissue surface with the syringe tip while simultaneously aspirating the resulting suspended tissue and mucus. To control for the significant amount of seawater and seawater-associated microbiota unavoidably captured during the collection of the slurry samples, a total of nine 10 ml syringes of ambient reef seawater were collected from approximately 1 m off the reef benthos, (hereafter referred to as “Syringe Method Control” samples). Immediately after collection, the syringes were placed in a Whirl-pak bag to prevent the loss of sample while underwater. Once back on board the boat, samples were transferred to 15 ml sterile conical tubes and placed in a 4°C cooler. Upon returning to the lab, samples were frozen to −20°C until analysis.

The physical and chemical environment of the surrounding seawater was characterized by measuring temperature, salinity, dissolved oxygen, pH, and turbidity using an Exo2 multiparameter sonde (YSI, Yellow Springs, OH, USA) (Table S1). The sonde probes were calibrated following manufacturer’s protocols on the day before sampling (February 10, 2020).

### 4.2 DNA extraction, PCR, and sequencing

Protocols for preparing samples for sequencing were specifically designed for the Illumina iSeq 100 System (Illumina Inc., San Diego, CA, USA), a portable, high-quality sequencing technology. In an approximately 1 cu. ft. size, the Illumina iSeq 100 System produces 4 million paired-end 150 bp sequence reads of high quality (<1% error rate) that can be offloaded and processed on a standard laptop without the need for Wi-Fi, making it an attractive technology to adapt for field-based microbiome studies. We brought the iSeq 100 System to a home rental in the USVI, which we transformed into a remote laboratory where we successfully conducted all DNA extractions, Polymerase Chain Reaction (PCR) and subsequent sequencing.

DNA was extracted from seawater, tissue and mucus slurry, and syringe method control samples, along with associated extraction controls, using the DNeasy PowerBiofilm Kit (Qiagen, Germantown, MD, USA). Modifications at the beginning of the extraction protocol were applied based on the sample type. For filtered seawater samples, the 0.22 μm filter was placed directly into the bead tube, and then manufacturer instructions were followed. For slurry and syringe method control samples, samples were thawed at room temperature, then immediately transferred to 4°C prior to extraction. Samples then were vortexed for 10 seconds and 1.8 ml of each sample was transferred to a bead tube. Samples were centrifuged at 12,045 rcf (maximum rcf available on centrifuge) for 10 min to concentrate tissue, mucus, and the associated microorganisms at the bottom of the tube, and supernatant was removed. For samples that were very clear (very little tissue collected via syringe) and for syringe method control samples, a second aliquot of 1.8 ml of sample was centrifuged on top of the existing pellet to capture more microorganisms. The extraction proceeded by following the manufacturer’s protocol. Six DNA extraction controls, three for each sample type, were generated by following the manufacturer’s protocol using: blank bead tubes for slurry and syringe method control samples (named D1-D3) and unused 0.22 μm filters placed in bead tubes for seawater samples (named D4-D6).

A two-stage PCR process was used to prepare the samples for sequencing. In the first stage, PCR was used to amplify the V4 region of the small sub-unit ribosomal RNA (SSU rRNA) gene of bacteria and archaea. The amount of DNA added and the total reaction volume of this first PCR varied by sample type. For each PCR, 2 μl of slurry and syringe method control template DNA was added to a final volume of 50 μl. 1 μl template in a 25 μl total reaction volume was used for seawater samples. For negative PCR controls, 1 or 2 μl of sterile PCR-grade water was used in 25 or 50 μl (total volume) reactions, respectively. One Human Microbiome Project mock community, Genomic DNA from Microbial Mock Community B (even, low concentration), v5.1L, for 16S rRNA Gene Sequencing, HM-782D was included as a sequencing control using 1 μl DNA in a 25 μl reaction. 50 μl reactions contained 0.5 μl polymerase (GoTaq, Promega, Madison, WI, USA), 1 μl each of 10 μM forward and reverse primers, 1 μl of 10 mM dNTPs (Promega), 5 μl MgCl_2_ (GoTaq), 10 μl 5X colorless flexi buffer (GoTaq), and 29.5 μl UV-sterilized, PCR-grade water. 25 μl reactions used the same proportions of reagents as 50 μl reactions. Earth microbiome project primers revised for marine microbiomes, 515F and 806R, targeted bacteria and archaea and were used with Illumina-specific adapters (Apprill *et al*., 2015; Parada *et al*., 2016). Two small, portable thermocyclers were used for the PCRs: the mini8 (miniPCR, Cambridge, MA, USA), which contained 8 wells and connected to a laptop for programming and initiation of the run, and the BentoLab (Bento Bioworks Ltd, London, UK), which contained 32 wells and was programmable as a unit. Using both machines was ideal because our targeted number of samples per iSeq 100 sequencing run was 40. The thermocycler program for the first stage PCR was: 2 min at 95°C, 35 cycles (coral slurry and syringe method control) or 28 cycles (seawater) of 20 sec at 95°C, 20 sec at 55°C, and 5 min at 72°C, followed by 10 min at 72°C and a final hold at 12°C. The final hold at 12°C was used due to the limitations of the BentoLab termocycler; samples were removed within an hour of the completed PCR program and stored at 4°C until purification. The resulting PCR products from slurry and syringe method control samples were purified as follows: 30 μl of PCR product per sample was mixed with 6 μl 5X loading dye (Bioline, London, UK) and separated using a 1.5% agarose gel stained with SYBR Safe DNA gel stain (Invitrogen, Thermo Fisher Scientific, Waltham, MA, USA). Bands of approximately 350 bp were excised by comparing to a 50 bp ladder (Bioline), and subsequently purified using the MinElute Gel Extraction Kit (Qiagen) following manufacturer protocols. For seawater PCR products, 5 μl of product mixed with 1 μl 5X loading dye was visualized on a 1% agarose gel to verify successful amplification, and the remaining PCR product was purified with the MinElute PCR Purification Kit (Qiagen).

The second stage PCR procedure attached unique index primers to each sample using the Nextera XT v2 set A kit (Illumina). Purified DNA (5 μl) from stage one PCR products was added to a 50 μl reaction with the following: 5 μl Nextera index primer 1, 5 μl Nextera index primer 2, 5 μl MgCl_2_ (GoTaq), 10 μl 5X colorless buffer (GoTaq), 0.5 μl Taq polymerase (GoTaq), 1 μl of 10 mM dNTPs (Promega), and 18.5 μl UV-sterilized, PCR-grade water. The PCR was run on the BentoLab or mini8 thermocyclers with the following program: 3 min at 95°C, 8 cycles of 30 sec at 95°C, 30 sec at 55°C, 30 sec at 72°C, followed by 5 min at 72°C and a final hold at 12°C. A subset of PCR products were visualized on a 1% agarose gel stained with SYBR Safe DNA gel stain (Invitrogen) using 5 μl product with 1 μl 5X loading dye (BioLine) to verify bands of approximately 450 bp, indicating successful attachment of sample-specific indexes. The stage two PCR products were purified with the MinElute PCR purification kit (Qiagen) following manufacturer protocols. Purified products were quantified using the Qubit 2.0 fluorometer dsDNA high sensitivity (HS) assay (Invitrogen) following manufacturer protocols to obtain stock concentrations in ng/μl. Concentrations were then converted to nM assuming average amplicon length of 450 bp and average nucleotide mass of 660 g/mol. Samples were diluted to 5 nM and pooled. Pooled samples were quantified via Qubit HS assay as before, and diluted to 1 nM, quantified again, and diluted to a loading concentration of 90 pM. A 10% spike-in of 90 pM PhiX Control v3 (Illumina, Inc.) was added to the pooled 90 pM library and 20 μl of the resulting library was run on the iSeq 100 System using paired-end 150 bp sequencing with adapter removal. Samples were sequenced over three sequencing runs.

### 4.3 Data analysis

All R scripts used for generating ASVs and producing figures were uploaded to GitHub. Forward reads were exclusively used for the downstream processing and data analysis due to minimal overlap between forward and reverse reads. The DADA2 pipeline (v.1.17.3; with parameters: *filterAndTrim* function: trimLeft = 19, truncLen = 145, maxN = 0, maxEE = 1, rm.phix = TRUE, compress = TRUE, multithread = TRUE) was used to remove the 515F and 806R primers from all sequence reads, filter the reads for quality and chimeras, and generate amplicon sequence variants (ASVs) for each sample (Callahan *et al*., 2016). This resulted in 17,190 ASVs of the same length (126 bp) across all samples. Taxonomy was assigned using the SILVA SSU rRNA database down to the species level where applicable (v.132) (Quast *et al*., 2012). ASVs that classified to mitochondria, chloroplast, eukaryote, or an unknown Kingdom were removed from the analysis, resulting in 7,366 remaining ASVs. We further filtered our dataset to remove possible contaminants introduced by DNA extraction reagents and introduced by seawater into coral tissue samples. The R package *decontam* (v. 1.6.0) was used to identify and remove DNA extraction contaminants in all samples (seawater, tissue/mucus slurry, and syringe method control) by using a combined frequency and prevalence method (Davis *et al*., 2018). The method identified 26 ASV contaminants, of which only 11 contained enriched frequency in DNA extraction controls so those 11 ASVs were removed (Appendix 1). Because the syringe method by nature collects a significant portion of seawater, the tissue/mucus slurry samples were, in essence, “contaminated” by seawater and thus, the signature of the ambient seawater microbiome needed to be removed from the slurry samples. To do this, the slurry samples were compared with the nine syringe method controls (seawater collected approximately a meter off of the benthos) using the prevalence method in *decontam*. The 184 ASVs identified as most prevalent in the nine syringe method controls (typically oligotrophic bacteria such as SAR11, *Prochlorococcus*, OM60 clade, *Synechococcus*, “*Candidatus* Actinomarina”, AEGEAN-169 clade, etc.) were generally found at low relative abundance in the tissue/mucus slurry samples (max relative abundance = 0.0074%) and were removed from the slurry sample ASV table (Appendix 2). After the ASVs identified as contaminants were removed, the tissue/mucus slurry samples and the near-coral seawater samples were re-merged into one large dataset. The re-merged dataset then was filtered to remove sparse ASVs (present at a count of 0 in the majority of samples) by removing ASVs with a count less than 0.5 when averaged across all samples. This left 2,010 ASVs, which were used for all downstream analyses.

Count data were transformed to relative abundance and coral tissue/mucus microbial communities were visualized using stacked bar charts. Data were then further log transformed following the addition of a pseudo count of one in preparation for beta diversity analyses. Bray-Curtis dissimilarity between samples was calculated using the R package *vegan* (v.2.5.7) and the resulting dissimilarities were presented in a Principal Coordinates Analysis (PCoA) (Oksanen *et al*., 2019). Permutational Analysis of Variance (PERMANOVA) with 999 permutations, using the *adonis* function in the *vegan* (Oksanen *et al*., 2019), compared the Bray-Curtis dissimilarity of healthy and diseased corals to test the hypothesis that coral microbiomes are significantly different between healthy and SCTLD-afflicted tissues. PERMANOVA was also used to test the hypotheses that species, reef location, and health state nested within species significantly structure the slurry microbial community. We tested the same hypotheses on the near-coral seawater directly overlying the coral colony to determine if species, reef location, or health drove microbiome community structure in near-coral seawater. Dispersion of beta diversity within coral tissue samples was calculated by measuring the distance to centroid within the PCoA as grouped by health state (HH and HD compared to DD) by implementing the *betadisper* function in *vegan* (Oksanen *et al*., 2019). Significant differences in dispersion by health state was determined by an independent Mann-Whitney U test. Additionally, variability of beta diversity was measured by extracting the Bray-Curtis dissimilarity values calculated within a tissue condition (diseased or healthy).

To detect ASVs enriched in diseased coral tissue compared to healthy tissue, the R package, *corncob* (v.0.1.0) (Martin *et al*., 2020), was employed, which modeled the relative abundance of each ASV and tested for differential abundance between healthy and diseased coral tissue. Following analysis of significantly differentially abundant ASVs in the coral tissue, we hypothesized that disease-associated ASVs would be recoverable in the near-coral seawater and graphed relative abundances of each disease-associated ASV in the near-coral seawater. We then employed corncob to test each identified disease-associated ASV to see if it was enriched at significantly higher abundances in seawater over diseased tissue compared to healthy tissue or apparently healthy colonies. Disease-associated ASVs were considered putative pathogens as they were enriched in diseased coral tissue/mucus slurries. Furthermore, we compared the ASV sequences of disease-associated ASVs to existing literature on SCTLD to determine if identical taxa were associated with SCTLD in other studies.

Sequences of putative pathogens, or disease-associated ASVs, were identified to the species level, when possible, as part of the DADA2 pipeline. To obtain better genus and species-level identification of putative pathogen ASVs and to relate these ASVs to other studies of coral disease-associated bacteria, we constructed phylogenetic trees for disease-associated ASVs classifying to *Vibrio, Arcobacter,* Rhizobiaceae, and Rhodobacteraceae. *Vibrio* and *Arcobacter* were targeted due to their increased representation in SCTLD-associated ASVs in this study and their previous association with SCTLD (Meyer *et al*., 2019) and coral disease in general (Ben-Haim *et al*., 2003; Ushijima *et al*., 2012). Rhizobiaceae and Rhodobacteraceae were targeted for phylogenetic tree analysis given their previous association with SCTLD (Rosales *et al*., 2020). Phylogenetic trees of coral-associated *Vibrio* and Rhodobacteraceae bacteria previously constructed from the Coral Microbiome Database (Huggett and Apprill, 2019) were used as reference trees for the insertion of SCTLD-associated ASVs that classified as *Vibrio* or Rhodobacteraceae. Insertion of our short SCTLD-associated sequence reads was achieved using the ‘quick add marked’ tool in ARB (version 6.0.6.rev15220) (Ludwig, 2004). Trees produced from ARB were exported using xFig. Phylogenetic trees for *Arcobacter* and Rhizobiaceae were constructed de novo using tools from the CIPRES Science Gateway (Miller *et al*., 2010). For each tree, long-read (∼1,200 bp) 16S rRNA gene sequences from closely-related (>90% sequence similarity) culture collection type strains, strains isolated from the marine environment, or clone sequences from corals were recovered via BLAST searches of SCTLD-associated ASVs from the present study or previous studies (Meyer *et al*., 2019; Rosales *et al*., 2020) to the non-redundant nucleotide collection, compiled into a FASTA file, and used for a sequence alignment in MAFFT (v7.402) (Katoh, 2002). This sequence alignment was then used to generate a reference tree using RAxML-HPC (v.8) (Stamatakis, 2014) with the following commands to produce a bootstrapped maximum-likelihood best tree: raxmlHPC-HYBRID -T 4 -f a -N autoMRE -n [output_name] -s [input_alignment] -m GTRGAMMA -p 12345 -x 12345. Next, SCTLD-associated short sequence reads were compiled into a FASTA file and added to the long-read sequence alignment in MAFFT using the “--addfragments” parameter. The sequence alignment with both short and long reads and the reference tree were then used as inputs for the Evolutionary Placement Algorithm, implemented in RAxML (Berger *et al*., 2011). RAxML was called as: raxmlHPC-PTHREADS -T 12 -f v -n [output_name] –s [long_and_short_read_alignment] -m GTRGAMMA -p 12345 -t [reference_tree]. The output tree including short read sequences (RAxML_labelledTree.[output_name]) was visualized and saved using the interactive tree of life (iTOL v5.6.3) (Letunic and Bork, 2016).

## Acknowledgements

The authors would like to thank Lei Ma, Joseph Townsend, Kathryn Cobleigh, Sonora Meiling, and Kelsey Beavers for field assistance and Illumina Inc. for technical assistance. Additionally, we would like to thank Laura Weber for assistance with iSeq protocol development and field assistance. This work was funded by The Tiffany & Co. Foundation, NSF OCE-1928761, and the Rockefeller Philanthropy Advisors, Dalio Foundation, and other generous donors of the Oceans 5 project. Samples were collected under permit DFW19057U. We would also like to thank Stefan Sievert, Harriet Alexander, and Tami Lieberman for helpful comments on the manuscript.

## Data Accessibility

Raw sequence reads were deposited into the NCBI GenBank under BioProject accession number PRJNA672912. Metadata associated with the study are also found at BCO-DMO under dataset 833133 (https://www.bco-dmo.org/dataset/833133). All R scripts used to generate figures and statistical tests are saved and publicly available on GitHub at https://github.com/CynthiaBecker/SCTLD-STT.

## Supplementary Figures and Tables

**Table S1.** Environmental conditions present at ‘Outbreak’ and ‘Existing’ diseased reefs.

| Reef | Outbreak | Existing |
| --- | --- | --- |
| Reef Name | Buck Island | Black Point |
| Lat (dd) | 18.27883 | 18.34450 |
| Lon (dd) | −64.89833 | −64.98595 |
| Depth (m) | 14.1 | 5.2 |
| Temperature (°C) | 26.88 | 26.98 |
| Salinity | 35.98 | 36.04 |
| Dissolved Oxygen (% sat.) | 102.5 | 105.0 |
| pH | 7.98 | 8.09 |
| Turbidity (NTU) | −0.05 | 0.12 |

**Fig. S1.**
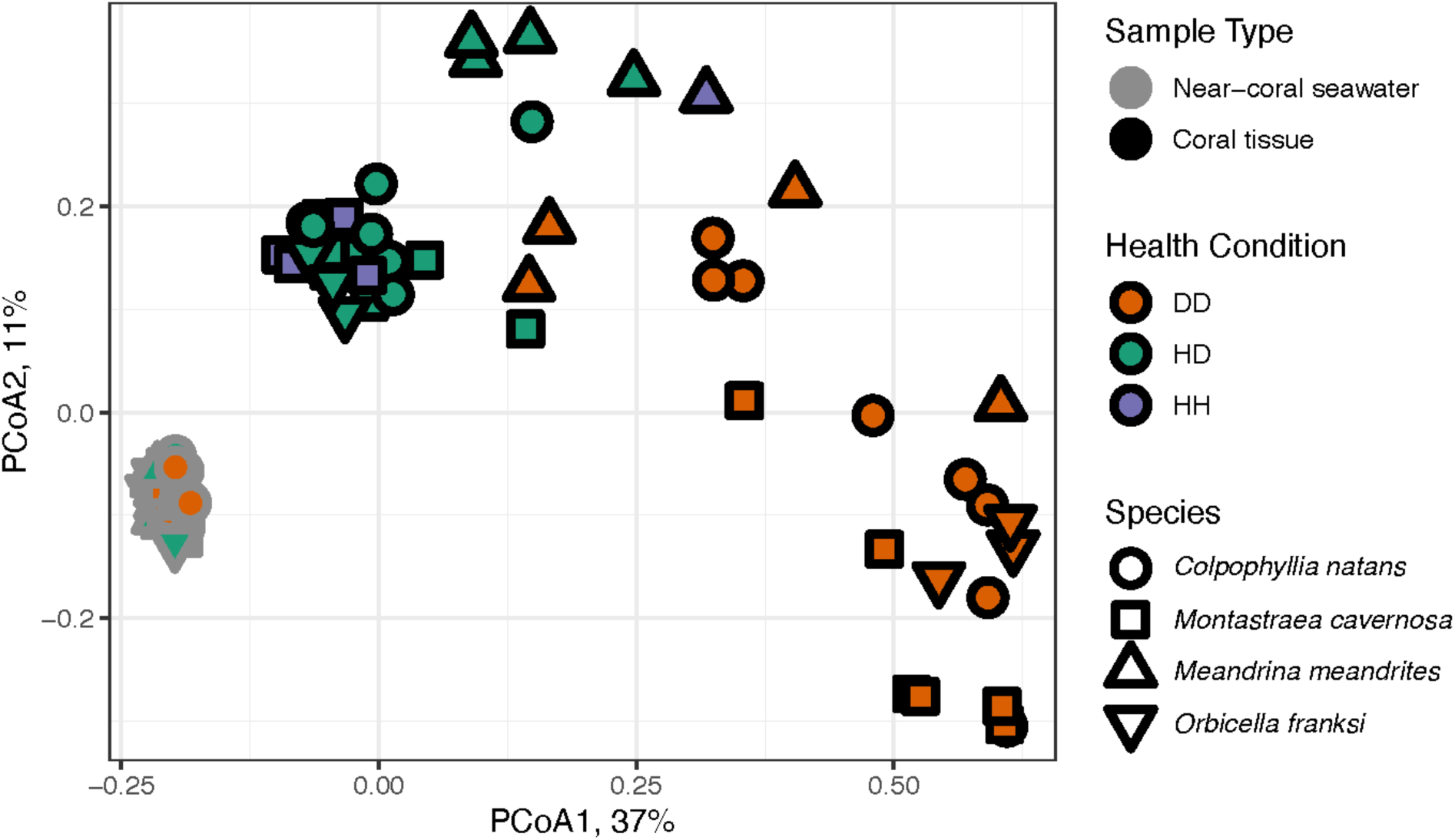
Principal coordinates analysis (PCoA) of Bray-Curtis dissimilarity between all coral and seawater samples. Seawater samples (gray outline) and coral tissue (black outline) samples are shaped by the coral species, *C. natans* (circle), *M. cavernosa* (square), *M. meandrites* (up triangle), and *O. franksi* (down triangle). Colors indicate health condition where DD = SCTLD lesion tissue, HD = healthy tissue on diseased colony, HH = healthy tissue from apparently healthy colony.

**Fig. S2.**
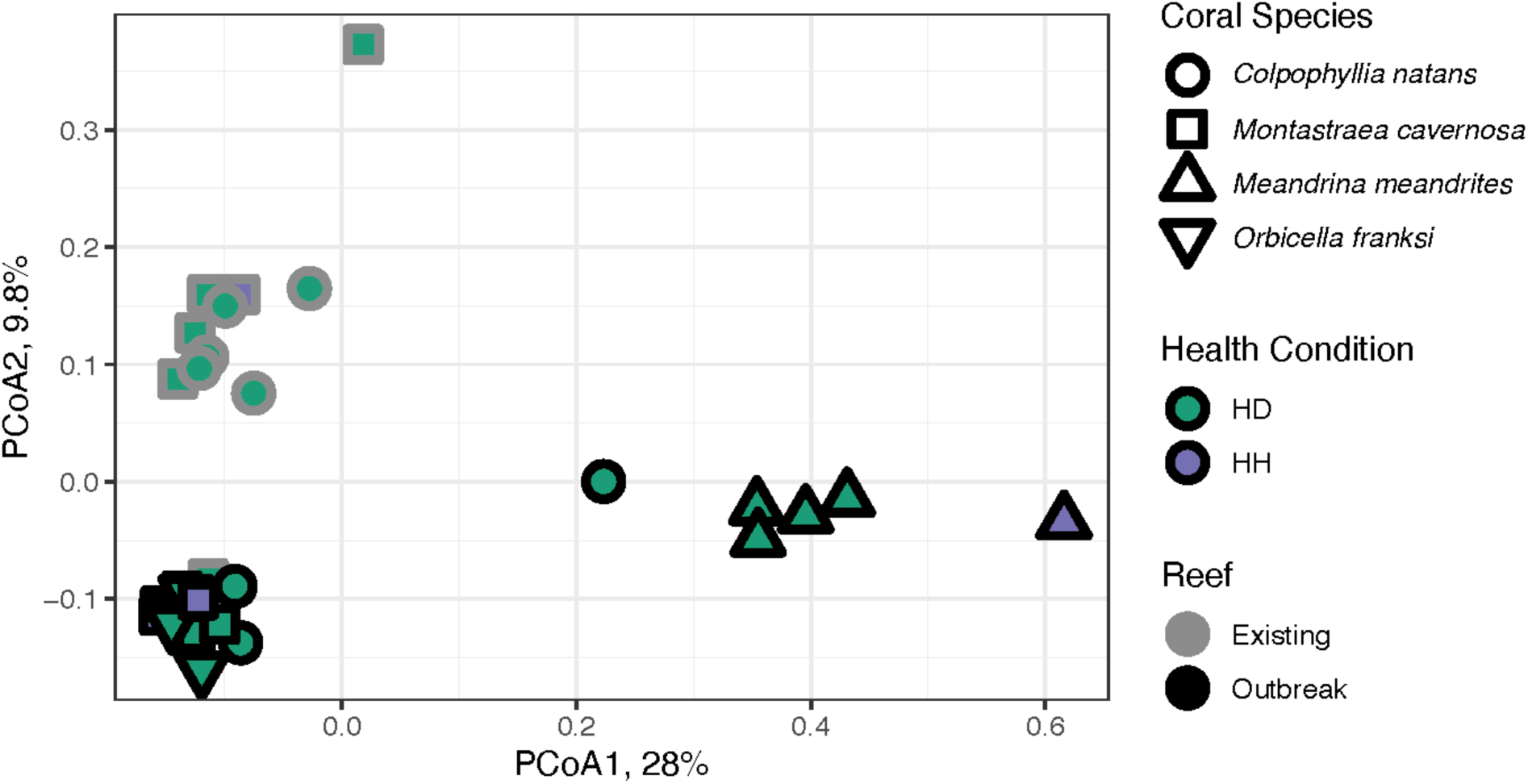
Principal coordinates analysis (PCoA) of Bray-Curtis dissimilarity within healthy coral tissue samples only. Outline color denotes whether the corals originated from the ‘Existing’ or ‘Outbreak’ SCTLD reef location. Fill color represents whether the healthy tissue sample was from a diseased colony (HD, green) or apparently healthy colony (HH, purple). Shape denotes the following coral species: *C. natans* (circle), *M. cavernosa* (square), *M. meandrites* (up triangle), and *O. franksi* (down triangle).

**Fig. S3.**
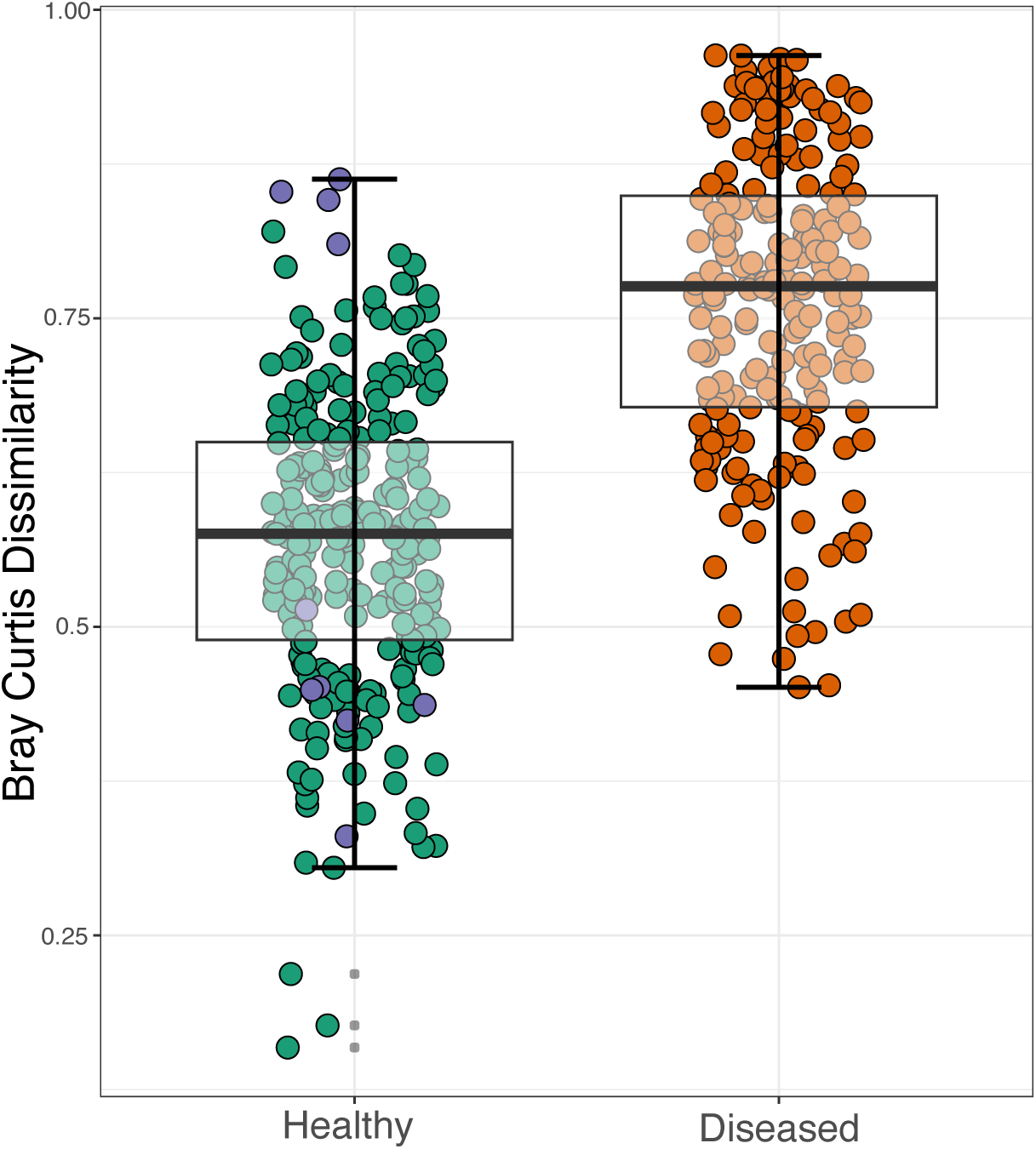
Boxplots denoting range in Bray-Curtis Dissimilarity values within healthy (HH = purple and HD = green) and diseased (DD = orange) coral tissue microbiomes. Difference between healthy and diseased Bray-Curtis dissimilarity values is significant by independent Mann-Whitney U Test (p < 0.001).

**Fig. S4.**
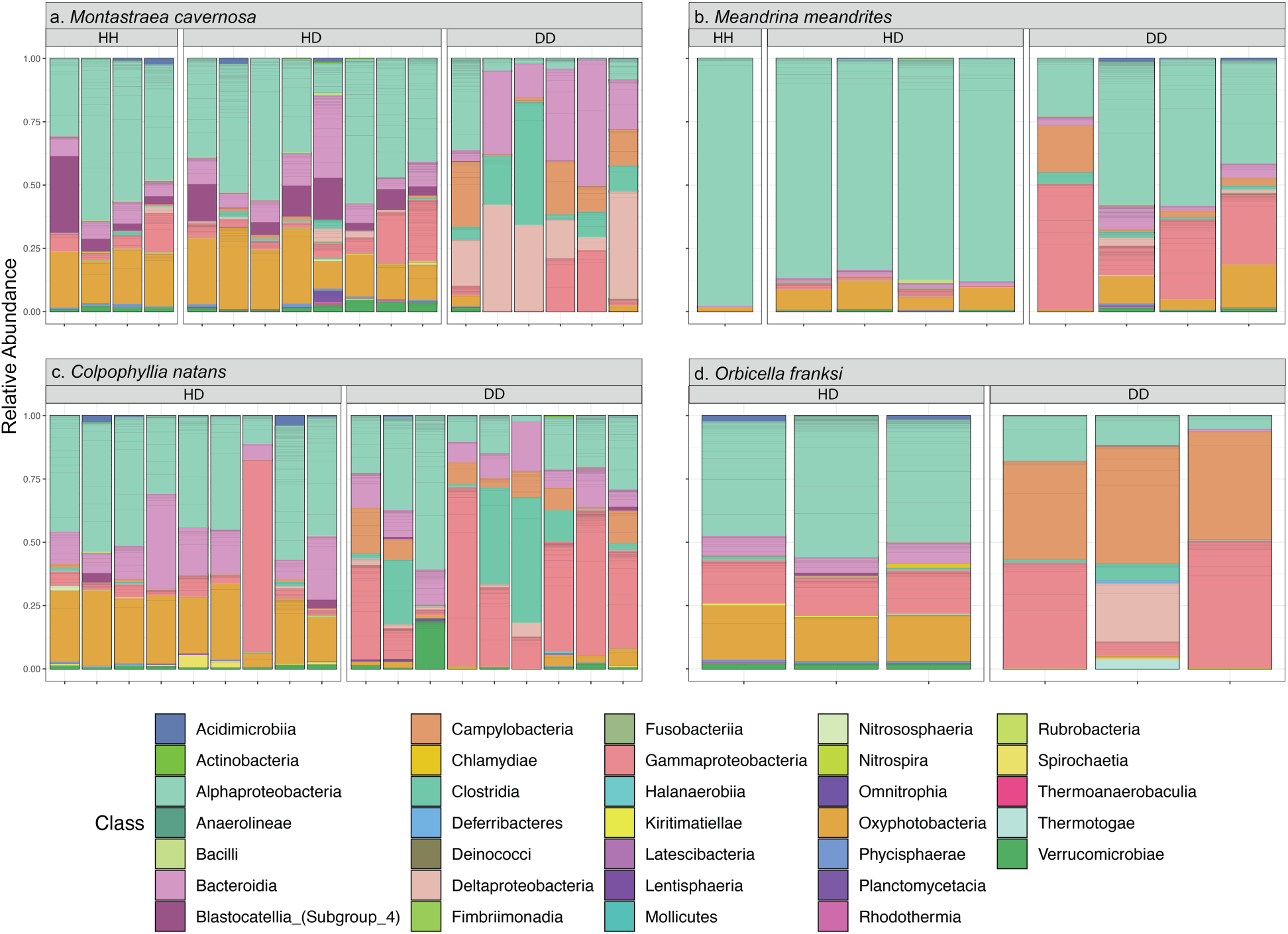
Stacked bar chart of microbial relative abundances within coral tissue in (a) *M. cavernosa*, (b) *M. meandrites*, (c) *C. natans*, and (d) *O. franksi*. Stacked bar charts are organized by tissue condition (HH = healthy tissue from a healthy colony, HD = Apparently healthy tissue from a diseased colony, DD = Disease lesion tissue from a diseased colony).

**Fig. S5.**
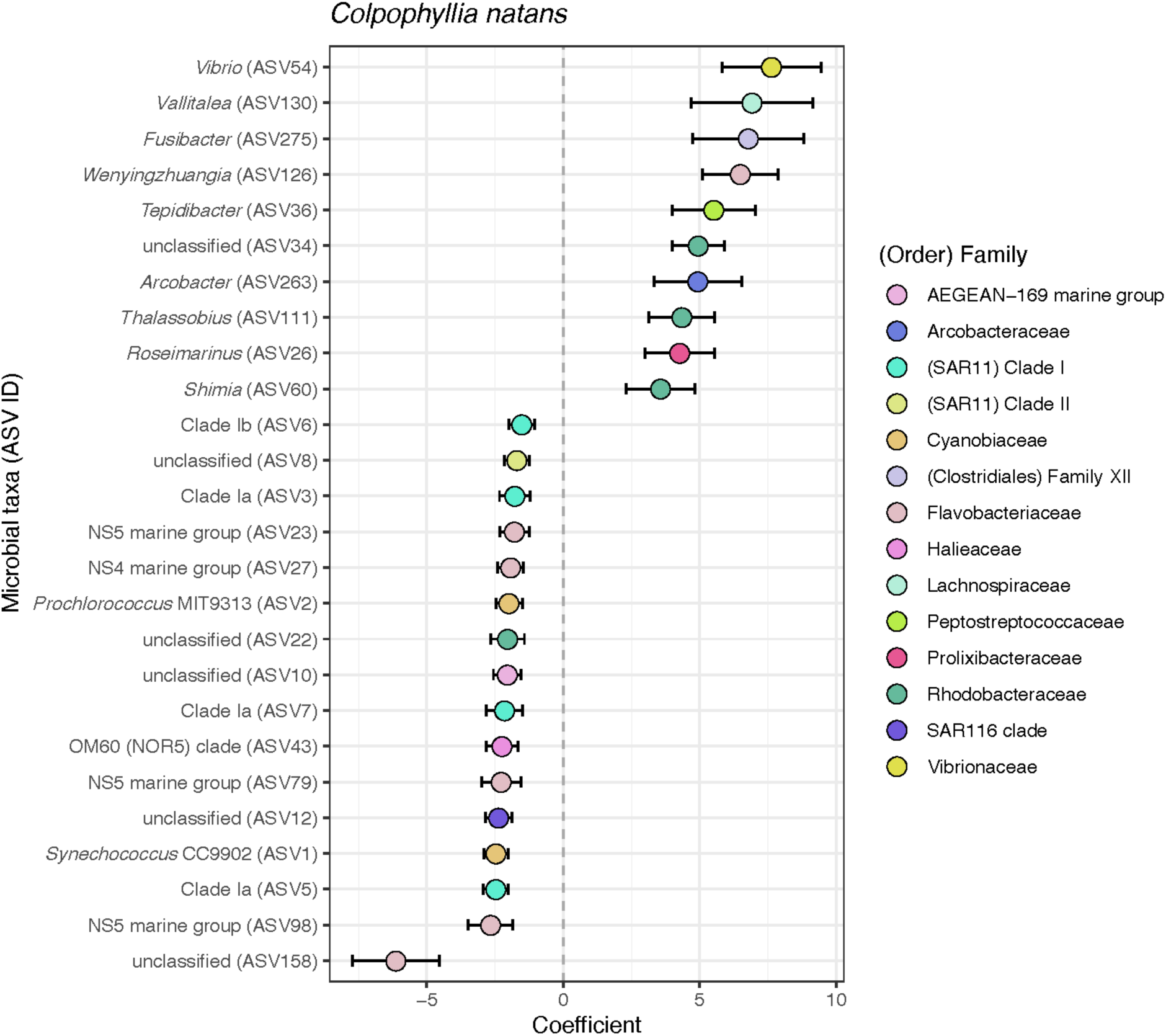
Significantly differentially abundant ASVs between diseased and healthy tissue in *Colpophyllia natans.* Positive coefficients indicate ASV relative abundance was enriched in diseased tissue relative to healthy tissue. Points are labeled by genera and ASV number, and colored by Family.

**Fig. S6.**
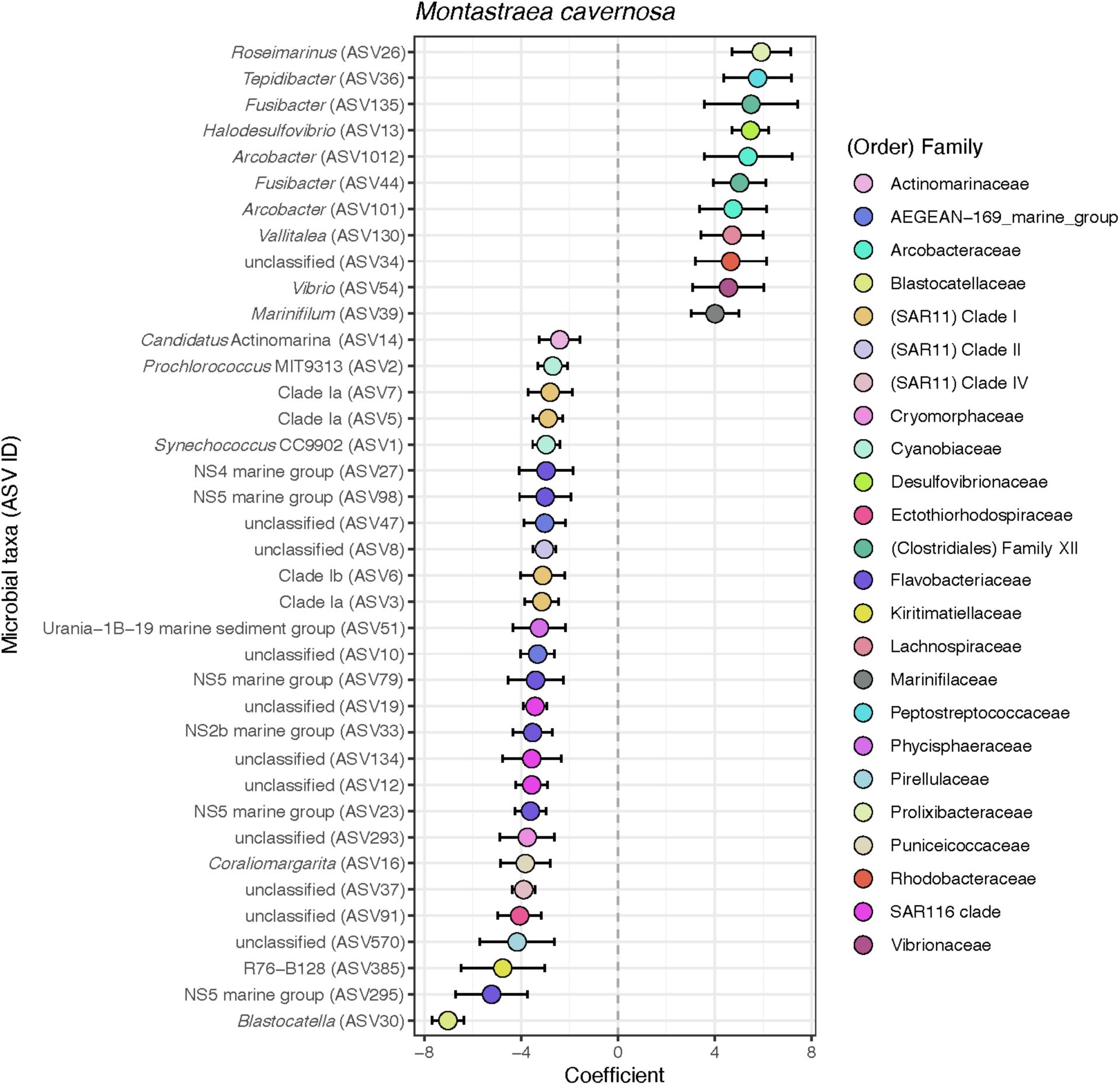
Significantly differentially abundant ASVs between diseased and healthy tissue in *Montastraea cavernosa.* Positive coefficients indicate ASV relative abundance was enriched in diseased tissue relative to healthy tissue. Points are labeled by genera and ASV number, and colored by Family.

**Fig. S7.**
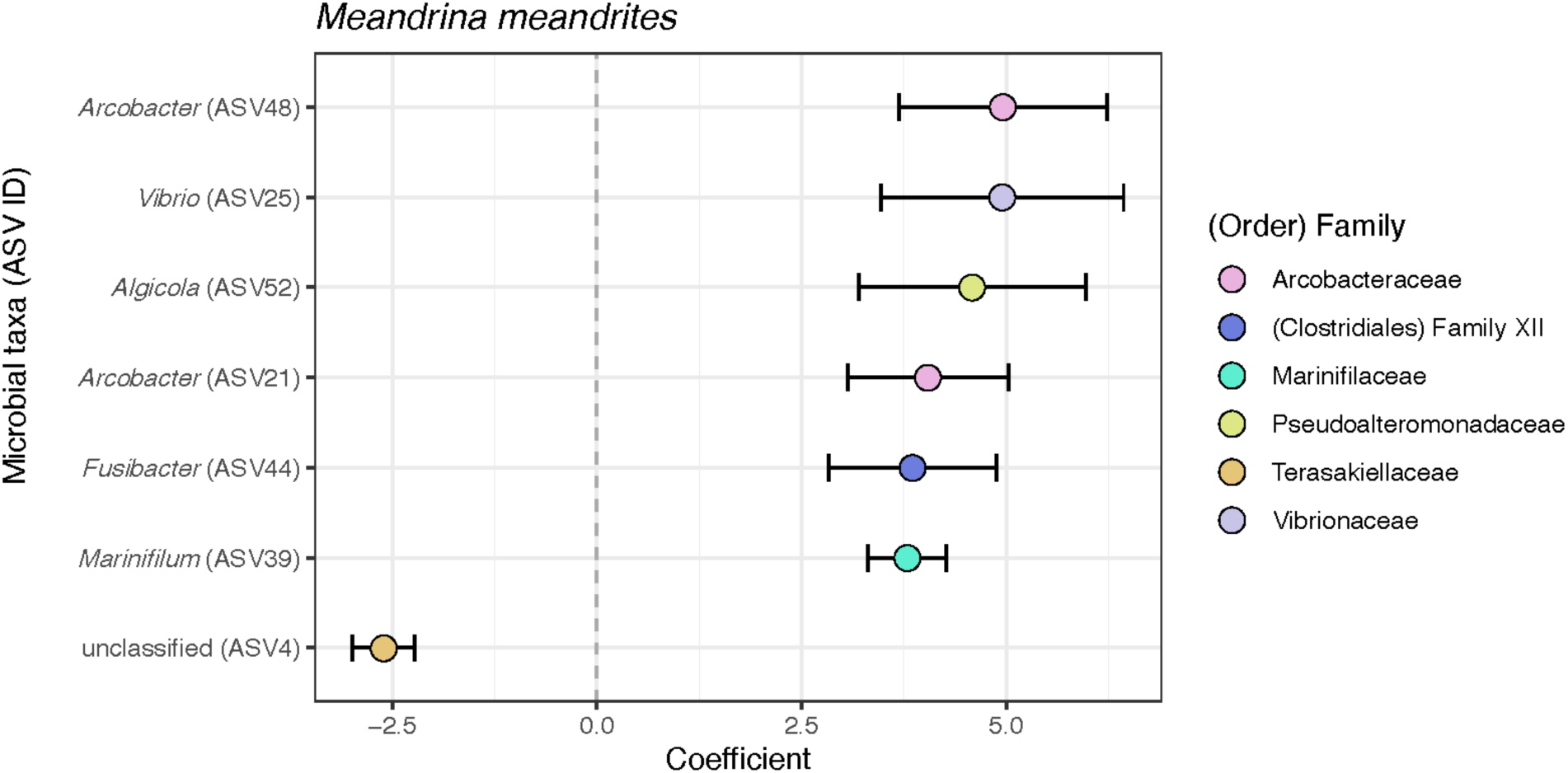
Significantly differentially abundant ASVs between diseased and healthy tissue in *Meandrina meandrites.* Positive coefficients indicate ASV relative abundance was enriched in diseased tissue relative to healthy tissue. Points are labeled by genera and ASV number, and colored by Family.

**Fig. S8.**
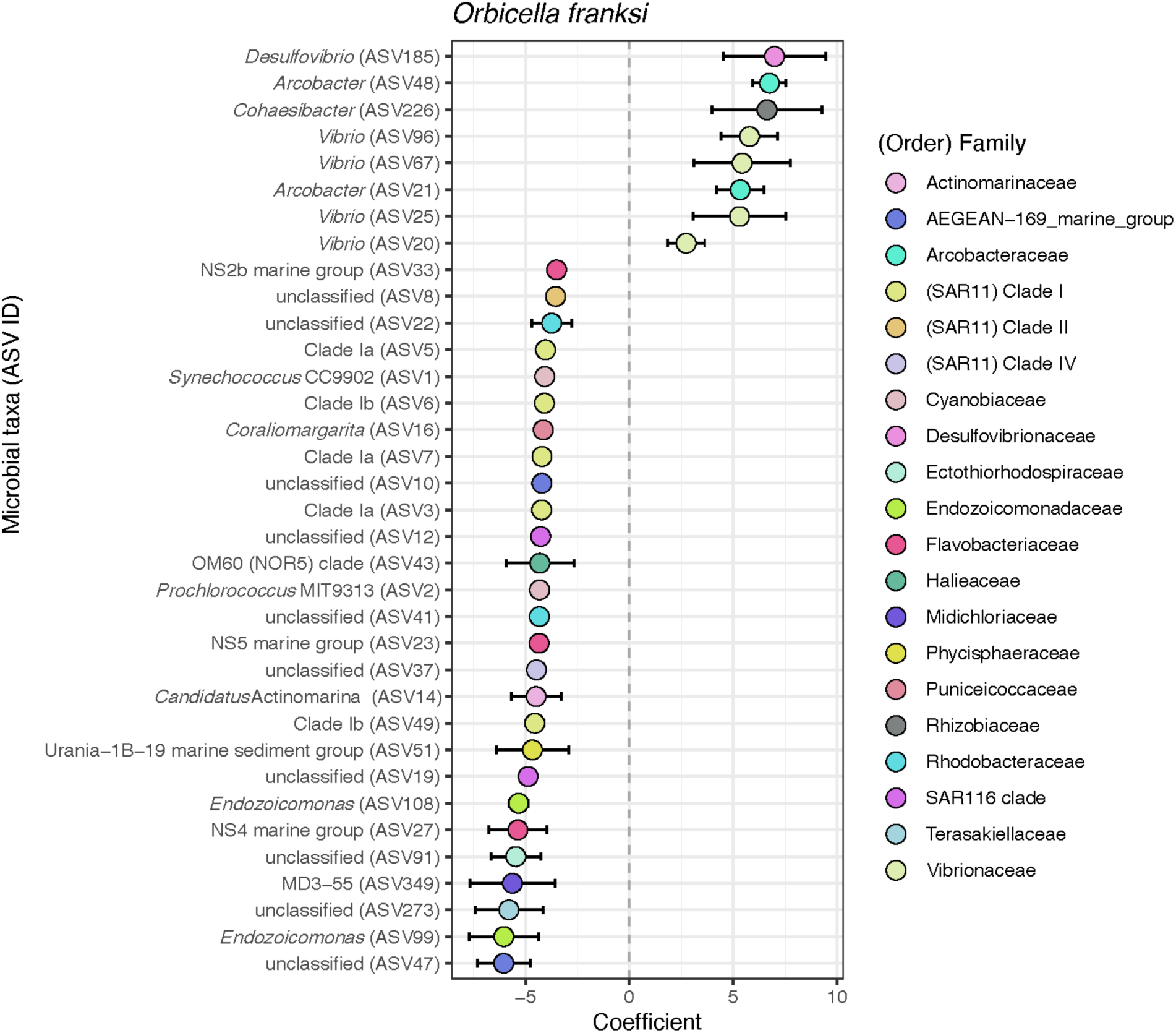
Significantly differentially abundant ASVs between diseased and healthy tissue in *Orbicella franksi.* Positive coefficients indicate ASV relative abundance was enriched in diseased tissue relative to healthy tissue. Points are labeled by genera and ASV number, and colored by Family.

